# Augmented prediction of multi-species protein–RNA interactions using evolutionary conservation of RNA-binding proteins

**DOI:** 10.1101/2025.09.27.678913

**Authors:** Jiale He, Tong Zhou, Lu-Feng Hu, Yuhua Jiao, Junhao Wang, Shengwen Yan, Siyao Jia, Qiuzhen Chen, Wentao Zhu, Jilin Zhang, Mutian Jia, Yuanning Li, Xianwei Wang, Yangming Wang, Yucheng T. Yang, Lei Sun

## Abstract

**Abstract:** RNA-binding proteins (RBPs) play critical roles in gene expression regulation. Recent studies have begun to detail the RNA recognition mechanisms of diverse RBPs. However, given the array of RBPs studied so far, it is implausible to experimentally profile RBP-binding peaks for hundreds of RBPs in multiple non-model organisms. Here, we introduce MuSIC (**<u>Mu</u>**lti-**<u>S</u>**pecies RBP–RNA **<u>I</u>**nteractions using **<u>C</u>**onservation), a deep learning-based framework for predicting cross-species RBP–RNA interactions by leveraging label smoothing and evolutionary conservation of RBPs across 11 diverse species ranging from human to yeast. MuSIC outperforms state-of-the-art computational methods, and provides predicted RBP-binding peaks across species with high accuracy. The prediction confidence is higher in the closely related species, partially due to the RBP conservation patterns. Finally, the effects of homologous genetic variants on RBP binding can be computationally quantified across species, followed by experimental validations. The target transcripts with disrupted binding events are enriched with the ubiquitination-associated pathways. To summarize, MuSIC provides a useful computational framework for predicting RBP–RNA interactions cross-species and quantifying the effects of genetic variants on RBP binding, offering novel insights into the RBP-mediated regulatory mechanisms implicated in human diseases.

**Highlights:**

- MuSIC integrates RBP-binding peaks with conservation-weighted label smoothing to predict RBP–RNA interactions across eleven species
- MuSIC outperforms state-of-the-art computational methods in predicting cross-species RBP–RNA interactions
- Cross-species prediction accuracy of RBP-binding peaks correlates with the conservation of RBPs
- MuSIC quantifies the effects of homologous SNVs on RBP binding with experimental validation in mouse

## Introduction

RNA-binding proteins (RBPs) play an important role that interact with intracellular RNAs by recognizing specific RNA motifs, and perform important biological functions including RNA transcription, splicing, transport, translation, and stability ^1–3^. Disruptions in RBP–RNA interactions have been implicated in the pathogenesis of various diseases, particularly cancer and neurodegenerative diseases ^4,5^.

Genetic variants, including single nucleotide polymorphisms (SNPs) and single nucleotide variants (SNVs), can alter the RBP-binding affinity, thereby disrupting RNA recognition and contributing to the development of various diseases ^6–8^. For example, the genetic variants in the iron-responsive element of *FTL* can impair the binding of IRP1 to *FTL* mRNA, leading to hyperferritinaemia-cataract syndrome ^9^. Similarly, the genetic variants in *TCF3* can hinder the binding of hnRNPH1 to exon 18b, thus disrupting alternative splicing of *TCF3* and promoting Burkitt lymphoma development^10^. These cases highlight the critical roles of RBP–RNA interactions in human disease pathogenesis.

Model organisms, such as mouse and zebrafish, are widely used in revealing the pathogenic mechanisms of complex human diseases^11,12^, where normal RBP–RNA interactions could be potentially disrupted by genetic variants ^13,14^. Various high-throughput experimental methods have been applied to identify RBP-binding peaks both *in vitro* ^15–17^ and *in vivo* ^18–21^. For example, PAR-CLIP technology has been applied to the mouse for exploring the regulatory roles of IGF2BP3 and LIN28B in hematopoietic reprogramming ^22^. However, it is still unrealistic to generate the binding profiles of hundreds of RBPs in human and animal models using *in vivo* high-throughput experimental methods.

With the recent advancements in AI technologies, computational methods leveraging large-scale RBP–RNA binding data can effectively learn RBP binding characteristics ^19,23,24^. Several computational methods using machine learning ^25,26^ and deep learning ^27–31^ technologies have been developed to predict RBP-binding peaks and model their RNA sequence preferences. However, the predictions by these traditional computational methods are generally restricted in the specific species, where the CLIP-seq datasets are generated. Given the fact that only a minority of the RBPs that have been studied by CLIP-seq technologies are from non-human organisms—65 RBPs identified in yeast, 45 in mouse, and only 5-6 in species such as fly, worm, and *A. thaliana* ^32^, it is vital to develop novel computational methods to predict RBP–RNA interactions in the cross-species tasks^33,34^.

The style transfer methods have been successfully applied in the field of computer vision, where similarity-guided label smoothing strategy could improve the cross-style performance ^35–37^. In our study, the analogies of the similarity features in label smoothing can be the sequence and structural conservation of RBPs. RBPs have been shown to be relatively highly conserved across species, particularly for the core RNA-binding domains (RBDs) ^38–40^. Consistently, the RBP-binding motifs from the RBPs containing highly conserved RBDs also show notable degree of conservation ^41–43^. These observations inspire us to apply the label smoothing strategy to the prediction of cross-species RBP–RNA interactions.

In this study, we develop MuSIC (**<u>Mu</u>**lti-**<u>S</u>**pecies RBP–RNA **<u>I</u>**nteractions using **<u>C</u>**onservation), a deep learning framework for cross-species RBP–RNA interaction prediction by leveraging the multi-level conservation patterns of RBPs. We show that MuSIC can accurately predict RBP-binding peaks across species and outperforms existing state-of-the-art computational methods, particularly in the closely related species. We then apply MuSIC to systematically profile the RBP-binding peaks and motifs for 186 RBPs across 11 species. Finally, we use MuSIC to quantify the effects of SNVs on RBP binding, providing insights into the RBP-mediated regulatory mechanisms associated with human diseases.

## Results

### Evolutionarily conserved RBPs exhibit similar RNA-binding preference

Previous studies have revealed that RBDs, the core functional elements in RBPs, exhibit a significantly slower rate of sequence evolution compared to genomic sequence context surrounding these RBDs, indicating their critical functions in regulating RNA recognition ^38^. We collected protein sequences and predicted structures of 216 RBPs from 11 species ranging from human to yeast, including human, orangutan, monkey, mouse, rat, chicken, frog, zebrafish, fly, *A. thaliana* and yeast, to systematically assess the conservation patterns of RBPs and their core RBDs (see **Methods** for details). Briefly, the conservation patterns of RBPs and their core RBDs was evaluated based on sequence and structure similarity, respectively (**Fig. 1a**, **Supplementary Fig. 1** and **Supplementary Table 1**). As expected, both sequence and structural similarities between human and non-human species declined with the increasing evolutionary distance (**Fig. 1a**). In particular, the protein sequence exhibited higher conservation for the core RBDs compared to the full-length RBPs; however, the structural similarity for both core RBDs and full-length RBPs remained relatively high across the 11 species ranging from human to yeast (**Fig. 1a** and **Supplementary Fig. 1**), indicating the fundamental importance of local structures of core RBDs for RNA recognition.

**Figure 1:**
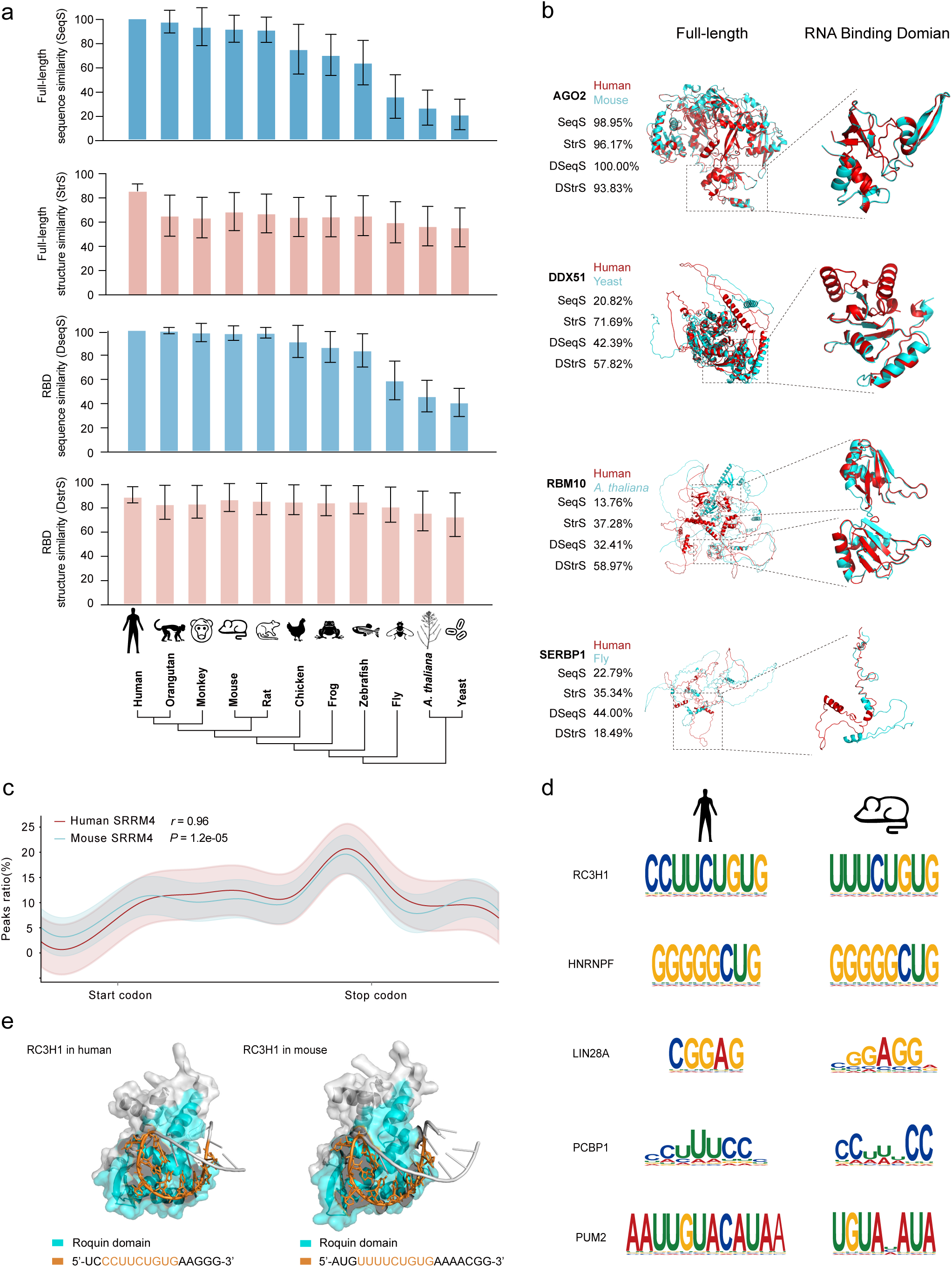
Evolutionary conservation of RBPs and their target RNAs. (a) Bar plots showing the sequence and structural similarity of RBPs across 11 species, in terms of full-length sequence (SeqS in blue), full-length structure (StrS in red), RBD sequence (DseqS in blue) and RBD structure (DStrS in red). The phylogenetic tree is constructed based on the 18S rRNA sequences in the 11 species, and shows the relative relationships between human and non-human species. (b) Examples of structural alignment of homologous RBPs, including AGO2 (the highly conserved), DDX51 (the considerable conserved), RBM10 (the intermediately conserved), and SERBP1 (the weakly conserved). The RBP structure in human (red) is compared to that in non-human species (blue). (c) SRRM4 binding distribution along meta-transcript in human (red line) and mouse (cyan line). (d) Binding motif alignment of homologous RBPs in human (left) and mouse (right). (e) Three-dimensional structure of RC3H1 in human (PDB ID 4QIL, left) and mouse (PDB ID 4QI2, right), showing bound RNA residues (orange), RBDs residues (blue), and other remaining RNA and protein residues (grey). See also **Supplementary Fig. 1** and **2**.

We then sought to partition the 216 RBPs into different groups based on their conservation patterns across the 11 species (**Supplementary Table 2**) and showed cases in each group (**Fig. 1b**): (i) highly conserved (1169, accounting for 49.0%), showing conservation in both sequence and structure in closely related species such as human and mouse, exemplified with the case of AGO2 protein; (ii) considerable conserved (769, accounting for 32.2%), showing conservation in full-length and RBD structure, such as DDX51; (iii) intermediately conserved (426, accounting for 17.8%), showing conservation in only RBD structure, such as RBM10; and (iv) weakly conserved (23, accounting for 1.0%), such as SERBP1. We found that almost all the RBPs (>99% of the total) exhibit structural conservation in their RBDs, consistent with previous studies showing that RBDs of the same RBP are highly structurally conserved across species ^44,45^.

Previous studies have shown that RBPs containing structurally conserved RBDs tend to recognize and bind their RNA target with similar sequence patterns ^41,45,46^. We first compared the distribution of RBP-binding peaks from the homologous RBPs along mRNAs in human and mouse, exemplified with the case of SRRM4 and LIN28A (**Fig. 1c** and **Supplementary Fig. 2a**). As expected, we observed a high correlation of the RBP-binding peak distributions between human and mouse (SRRM4 r = 0.96, LIN28A r = 0.91, Spearman’s rank correlation test). Notably, SRRM4 showed stronger preference of binding on the stop codon and a depletion around the start codon (**Fig. 1c**), suggesting the critical roles of SRRM4 in regulating alternative splicing of RNA within these regions ^47^. Next, we explored the motif-level conservation of RBPs using the ATtRACT database ^48^, which contains experimentally validated binding motifs in multiple species. Comparative analysis of RBP-binding motifs between human and mouse revealed a high degree of conservation (e.g., UUCUGUG motif for RC3H1, GGGGGCUG motif for HNRNPF, GGAG motif for LIN28A) (**Fig. 1d**). Moreover, both the structures derived from X-ray diffraction (**Fig. 1e**) and AlphaFold3 ^49^ prediction (**Supplementary Fig. 2b-e**) suggest that the homologous RBPs from different species exhibit similar three-dimensional interaction patterns between the interface residues in RBPs and their recognized target binding motifs (PDB ID 4QIL in human and 4QI2 in mouse) (**Fig. 1e**). Taken together, these results suggest that RBPs show high sequence and structural conservation across species, and maintain high conservation in their RNA-binding patterns, including motifs, binding regions within transcripts, and key interacting residues.

Inspired by the aforementioned observations and previous studies^30,33,43^, we aim to develop a novel computational model to systematically predict the RBP–RNA interactions in the target species (e.g., non-human species) by integrating the RBP-binding peaks from the source species (e.g., human) and the RBP similarity patterns between species. We leveraged the large-scale RBP-binding peaks from different species (human, mouse, and zebrafish) to construct the computational model. We initially found 23 homologous RBPs between human and mouse, and one homologous RBP between human and zebrafish with available datasets from POSTAR3 (**Supplementary Fig. 2f**), among which 38 datasets from 14 RBPs were curated for the training and validation of the cross-species model (see **Methods** for details; **Supplementary Table 3**).

### Overview of MuSIC algorithm

We developed MuSIC, a deep learning framework that could integrate RBP-binding peaks and cross-species conservation information of RBPs to accurately model and predict RBP–RNA interactions in diverse non-human species. Briefly, the MuSIC framework consists of four components (**Fig. 2a**): (i) pre-processing input data: collecting peaks of RBPs from multiple species, and constructing training and validation sets; (ii) feature engineering: incorporating nucleotide sequences and RNA secondary structures to learn RBP-binding preferences and enhance the prediction ability of the model (**Fig. 2b**); (iii) predicting RBP–RNA interactions: dynamic optimization on the prediction results using label smoothing regularization and gradient weight adaptation in a convolutional neural network (CNN); and (iv) analyzing RBP– RNA interactions: cross-species motif identification and other downstream applications.

**Figure 2:**
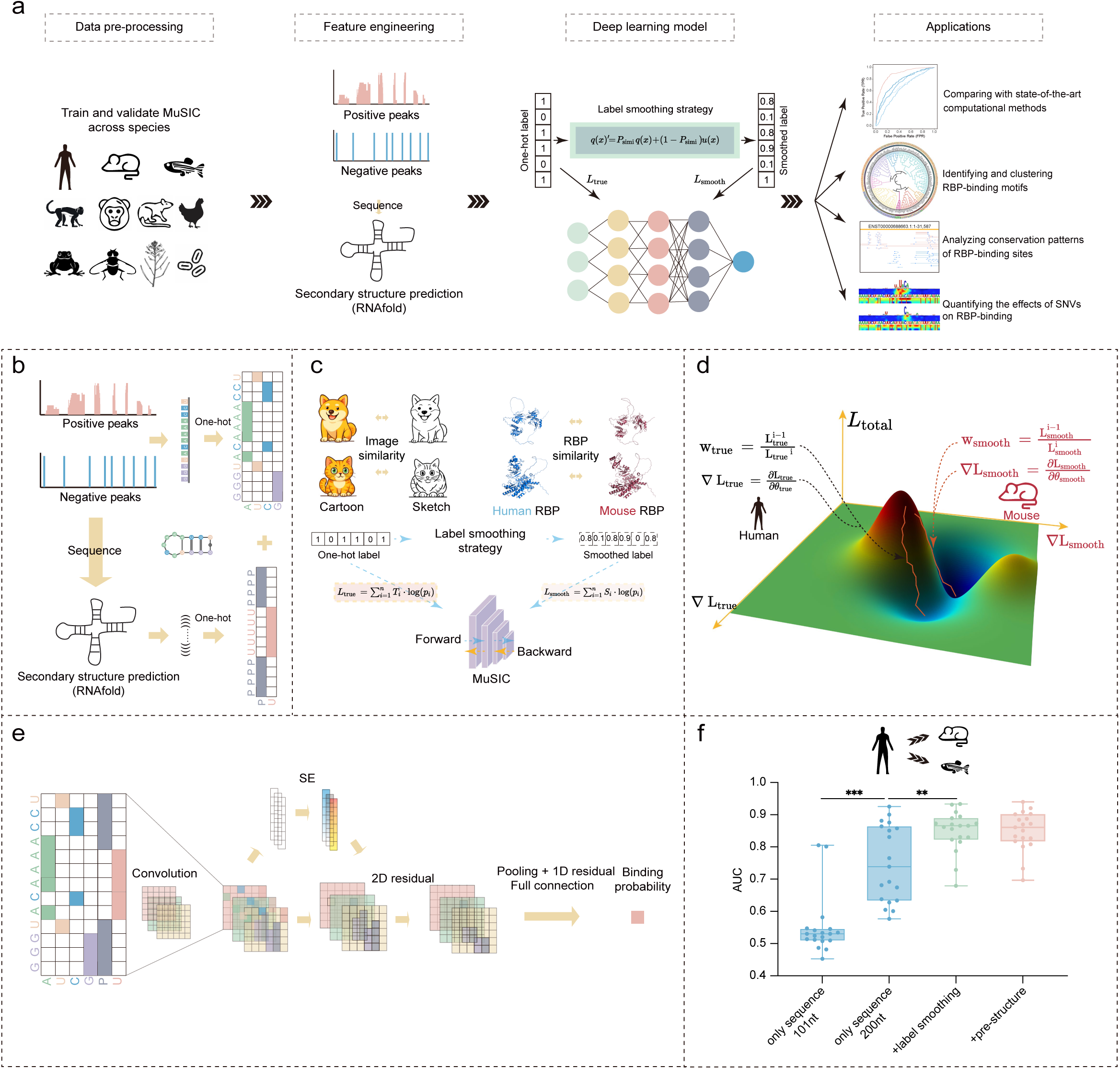
The workflow of MuSIC. (a) Schematic showing the workflow of MuSIC. Step 1 (data pre-processing): collecting peaks of RBPs from multiple species, defining the positive and negative peaks, and constructing the training and validation set. Step 2 (feature engineering): encoding for the RBP-binding peaks and predicted RNA structures. Step 3 (Deep learning model): predicting RBP–RNA interactions using a CNN framework integrated with label smoothing strategy. Step 4 (Applications): comparing prediction performance with state-of-the-art computational methods, identifying and clustering RBP-binding motifs, analyzing conservation patterns of RBP-binding peaks, and quantifying the effects of SNVs on RBP binding. (b) Flowchart showing data pre-processing and feature encoding. Top: positive (red) and negative (blue) peaks are encoded into one-hot vectors. Bottom: computationally predicted RNA structures are encoded into one-hot vectors. (c) Diagram showing label smoothing strategy. Top left: using the image similarities between Cartoon and Sketch in computer vision for label smoothing strategy. Top right: using the RBP similarities between human (blue) and mouse (red) for label smoothing. Bottom: transforming the one-hot labels into smoothed labels based on RBP conservation and computing cross-entropy loss for both one-hot labels and smoothed labels. (d) Diagram showing two components of the total loss *L*_total_, *L*_*ture*_ (human in black) and *L*_*smooth*_ (mouse in red). (e) Overview of the deep learning model architecture. One-hot encoded input (b) is processed sequentially through the convolutional layers, SE modules, residual blocks, max pooling, and fully connected layers to predict the binding probability. (f) Boxplot showing the prediction performance for different types of input features, including only sequence (101nt / 200nt) (blue), 200nt sequence with label smoothing (green), sequence with label smoothing and predicted structure (red). See also **Supplementary Fig. 3**.

We realized that the cross-species prediction tasks often have great challenges in generalization, e.g., the ability to accurately predict RBP–RNA interactions in mouse based on the same distribution as the training data in human ^33,34^. There are several disadvantages in the existing deep learning-based methods, such as overfitting caused by the limitations of one-hot label loss functions, and heightened sensitivity to label noise and boundary ambiguities inherent in one-hot supervision ^36^. To this end, we leveraged the label smoothing strategy that has been widely used in the field of computer vision to facilitate the cross-style classification ^35^. In MuSIC, the conservation of RBPs across species can be used to enhance the model generalization and mitigate overfitting in a similar way. Briefly, MuSIC employs a novel approach in applying the label smoothing strategy, in which the one-hot labels were converted into soft labels based on the pre-defined RBP conservation scores ^37^. The cross-entropy loss was computed for both label types, with the final loss obtained by weighting and summing the two components (**Fig. 2c**). In addition, we used gradient weight adaptation algorithm to dynamically adjust the weight of each loss component during the model training (**Fig. 2d**). Next, the MuSIC model combines several convolutional layers, residual blocks and the Squeeze-and-Excitation (SE) module to capture and enhance RBP binding features learning, in which overfitting could be minimized by dropout and early stopping (**Fig. 2a, e**). Finally, the learned features were integrated using the fully connected layers to generate the prediction outputs (see **Methods** for details).

Existing state-of-the-art computational methods typically utilize input sequence lengths of 101nt ^28,30,31^ or 400nt ^26,50^. We found that increasing the input sequence length to 200nt could significantly improve the Area Under the Curve (AUC) score in cross-species prediction (**Fig. 2f** and **Supplementary Fig. 3a**). The inclusion of label smoothing, and gradient adaptation could further improve the AUC score (**Fig. 2f** and **Supplementary Fig. 3b**). Notably, incorporating predicted RNA structures could further enhance the AUC score to 0.85 in cross-species prediction (**Fig. 2f**). We also evaluated the prediction performance using the experimentally measured *in vivo* RNA structures instead of the predicted RNA structures (**Supplementary Fig. 3c**). The predicted RNA structures were used in the model because of the limited performance gains by using the *in vivo* RNA structures, as well as very few *in vivo* RNA structure datasets are available in other species. Taken together, these results reveal that extending the sequence length and applying label smoothing could significantly improve the performance of cross-species generalization.

We then characterized the contribution of each component of the MuSIC framework to the performance of feature learning. The positive and negative samples of input RNA sequences cannot be clearly separated in the original feature space using t-SNE (**Supplementary Fig. 3d**). After adding the SE module and residual block processing, the t-SNE clustering performance on the positive and negative samples could be significantly improved, thus leading to the apparent discriminative power of the final output from the fully connected layers (**Supplementary Fig. 3d**).

The classification performance of MuSIC’s classifier was benchmarked against diverse machine learning and deep learning classifiers using the identical input features. We found that MuSIC consistently outperformed all the baseline classifiers, confirming the robustness and effectiveness of MuSIC in modeling RBP binding across species (**Supplementary Fig. 3e, f**).

### MuSIC outperforms existing computational methods in predicting cross-species RBP–RNA interactions

We evaluated the prediction performance of MuSIC using 262 datasets from 186 RBPs, benchmarked under both within-species (224 datasets from 186 RBPs) and cross-species (38 datasets from 14 RBPs) scenarios (**Supplementary Table 3**). All the datasets were processed using a uniform preprocessing pipeline to ensure fairness in the performance evaluation (see **Methods** for details). Briefly, we performed a systematic comparison of prediction accuracy with four state-of-the-art computational methods, including HDRNet ^31^, PrismNet ^30^, DeepBind ^28^, and GraphProt ^26^. As expected, MuSIC showed better prediction accuracy than other methods for the within-species prediction (AUC: MuSIC = 0.88, HDRNet = 0.87, PrismNet = 0.86, DeepBind = 0.67, GraphProt = 0.72) (**Supplementary Fig. 4a**).

We then evaluated the prediction accuracy of MuSIC and other computational methods in the cross-species prediction, where we trained the model in human and validated in mouse and zebrafish (19 datasets). MuSIC outperformed other methods, showing an obvious improvement of in AUC (AUC: MuSIC = 0.85, HDRNet = 0.73, PrismNet = 0.73, DeepBind = 0.71, GraphProt = 0.70; **Fig. 3a**), as well as in the Area Under the Precision-Recall Curve (PRAUC) and accuracy (ACC) metrics (**Supplementary Fig. 4b, c**). In addition, we found that MuSIC a significant performance improvement of AUC for predicting RBP–RNA interactions in the closely related predictions, e.g., trained in human and validated in mouse (**Supplementary Fig. 4d-f**). For example, MuSIC showed AUCs of 0.88 for YTHDC2 and 0.89 for TAF15 in the cross-species prediction, a significant improvement than other computational methods (**Fig. 3b** and **Supplementary Fig. 5a**). Taken together, these results showed that MuSIC could improve the prediction accuracy of RBP–RNA interactions than other computational methods, especially under the cross-species scenario. Consistently, MuSIC showed better performance in separating the positive and negative samples of input RNA sequences in the feature space than other computational methods (**Fig. 3c** and **Supplementary Fig. 5b**).

**Figure 3:**
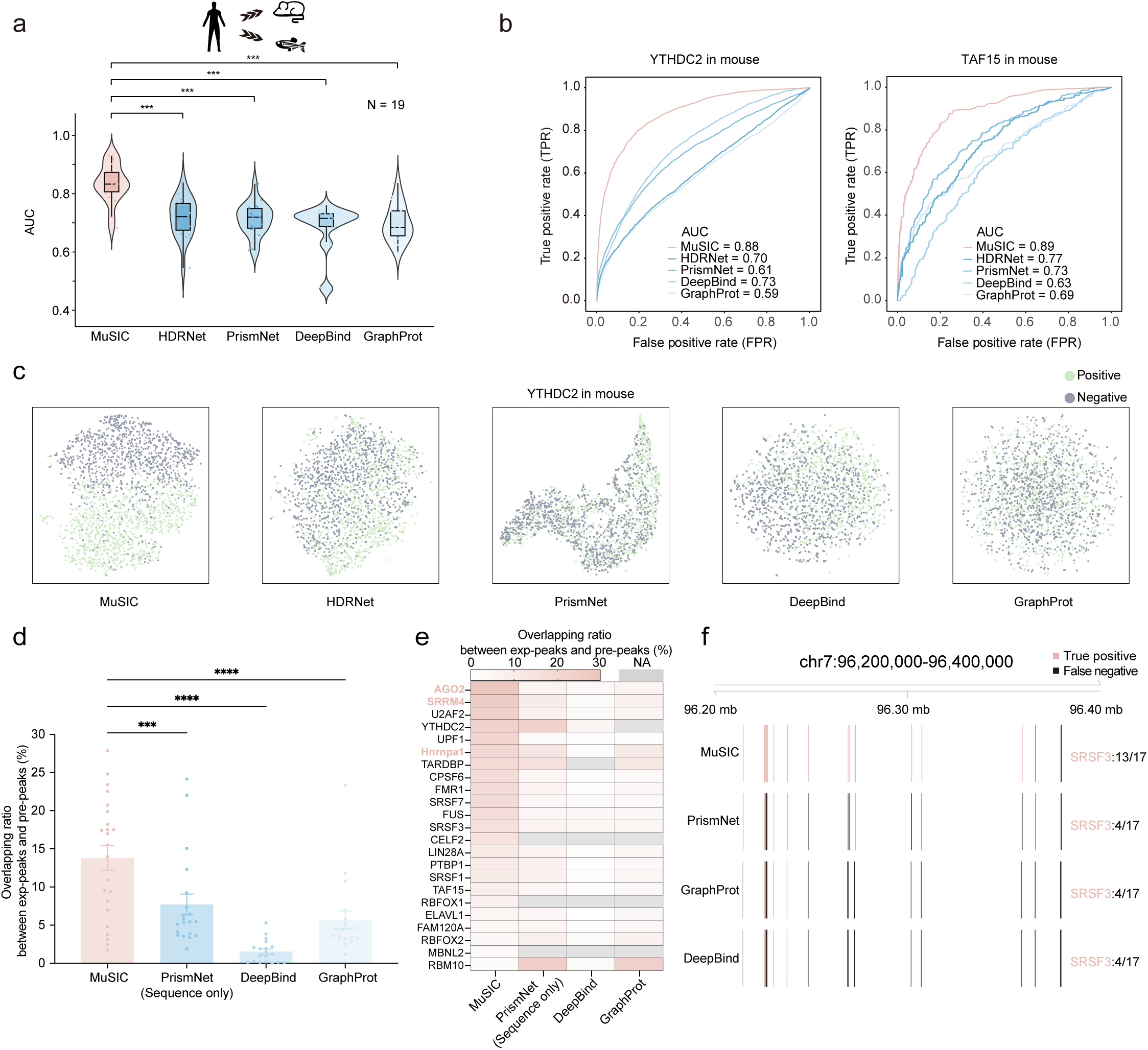
Comparison of prediction performance between MuSIC and other computational methods. (a) Violin plot showing the prediction accuracy of MuSIC (red) and other computational methods (blue) for 19 cross-species datasets. *** *P* < 0.005, paired paired t-test. (b) ROC curves showing the prediction accuracy of MuSIC (red) and other computational methods (blue) for two RBPs, YTHDC2 (left) and TAF15 (right). (c) t-SNE clustering showing the output feature maps by MuSIC and other computational methods for separating the positive (cyan) and negative (grey) peaks. (d) Bar plot showing the overlapping ratio between the experimentally-derived and predicted peaks by MuSIC (red) and other computational methods (blue). *** *P* < 0.005, paired t-test; ****, *P* < 0.0001, paired t-test. (e) Heatmap showing the overlapping ratio between the experimentally-derived and predicted peaks by MuSIC and other computational methods. (f) An example of predicted peaks distribution along the local region in mouse genome (chr7:96,200,000–96,400,000). Red indicates true positives and black indicates false negatives, as evaluated against the experimentally-derived peaks. See also **Supplementary Fig. 4, 5** and **6**.

Finally, we compared the prediction accuracy of MuSIC and other computational methods by evaluating to what degree the RBP-binding peaks from CLIP-seq experiments could be rediscovered (see **Methods** for details). Briefly, the model was trained based on the RBP-binding peaks in human, followed by predicting peaks across all the transcripts in mouse (for 22 RBPs) and zebrafish (for 1 RBP) (**Supplementary Table 4**). For each RBP, we observed that MuSIC showed consistently better performance than other computational methods (**Fig. 3d**), particularly on the cases of AGO2, SRRM4, and Hnrnpa1 (**Fig. 3e**). Notably, HDRNet was excluded in this analysis due to its reliance on the experimental RNA structure features, which were not available in the relevant mouse datasets. For all the 23 RBPs analyzed here, the predicted peaks by MuSIC showed high correlation with the experimentally derived ones (r = 0.73, Spearman’s rank correlation test; **Supplementary Fig. 6a-c**). Here, we showcased the site-level prediction performance of MuSIC using SRSF3 within the local genomic region (chr7: 96,200,000–96,400,000) in mouse, where 13 out of 17 experimentally derived peaks could be successfully recovered by MuSIC (**Fig. 3f**). In contrast, only 4 peaks could be predicted by other computational methods. These findings confirm the feasibility and effectiveness of MuSIC in accurately capturing RBP–RNA interactions across species.

### MuSIC generates systematic RBP-binding predictions in diverse species

The aforementioned results reveal that MuSIC could generate better cross-species predictions than other computational methods, particularly when the source and target species are more evolutionarily closed. It will be useful to explore to what degree the source and target species are evolutionarily closed, then the MuSIC can generate reliable prediction results. We then evaluated the relationship between the prediction accuracy and the RBP conservation between species in terms of both their sequence and structure (**Supplementary Fig. 7a**; see **Methods** for details). Notably, we observed a strong positive correlation (r = 0.76, Pearson’s correlation test; **Fig. 4a**) between the prediction accuracy (measured by AUC) and the RBP conservation based on 14 RBPs shared between mouse and zebrafish, indicating that the degree of RBP conservation could be used to quantificationally threshold the prediction results of RBP–RNA interactions. We then predicted the RBP–RNA interactions for the 10 species in a broad evolutionary range using MuSIC, and filtered low-confidence prediction results based on the degree of RBP conservation equivalent to AUC of 0.8 (see **Methods** for details).

**Figure 4:**
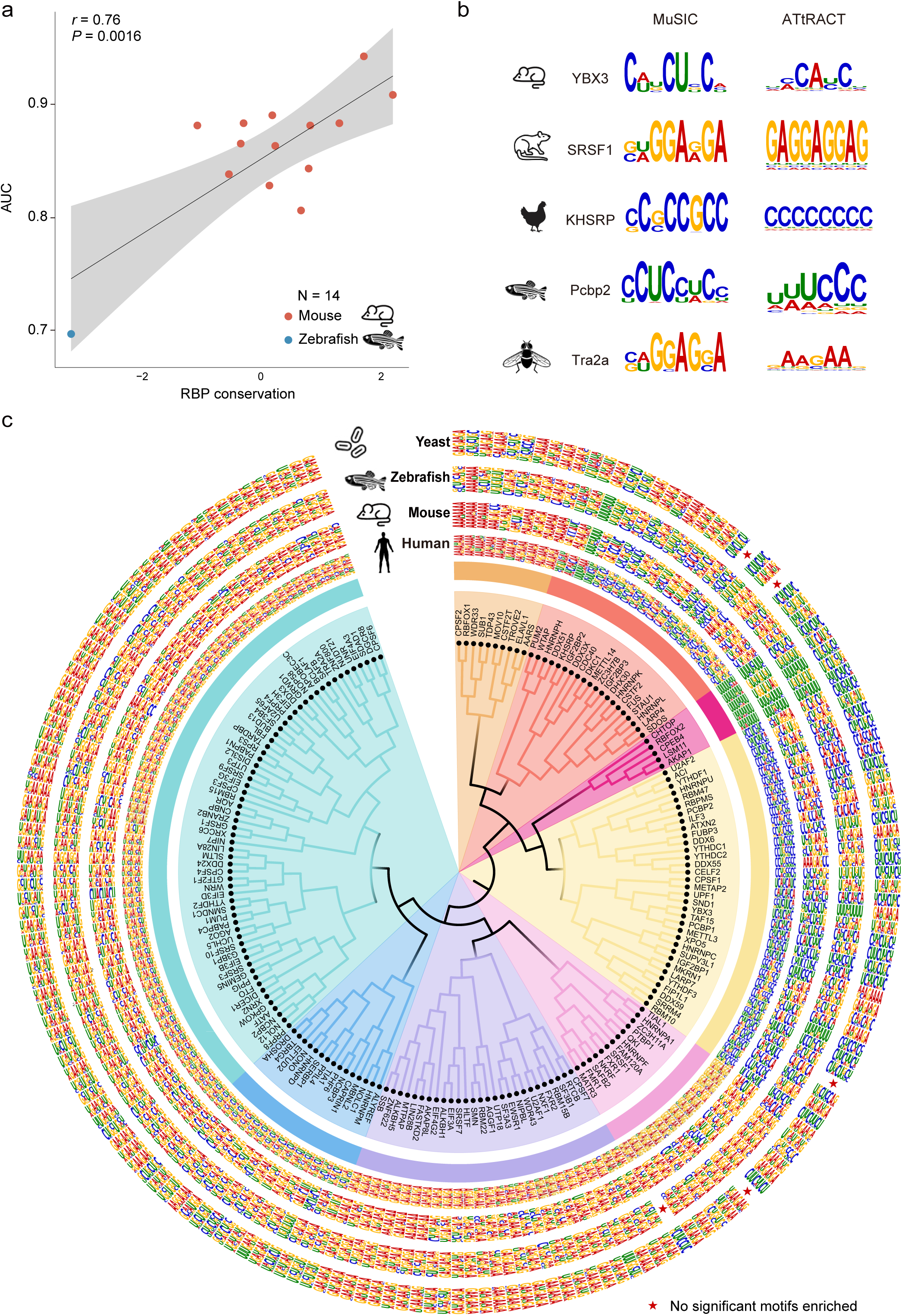
Reliable predictions of RBP-binding peaks and motif clustering by MuSIC. (a) Scatter plot showing the correlation between the cross-species prediction accuracy and RBP conservation for the 14 RBPs (mouse in red, zebrafish in blue). (b) Examples showing the consistency of the predicted binding motifs and experimentally-derived motifs (from ATtRACT database) of several RBPs. (c) Hierarchical clustering of 186 predicted binding motifs in human, mouse, zebrafish, and yeast. Stars indicate that there are no motifs enriched. See also **Supplementary Fig. 7** and **8**.

As expected, more than 90% of the RBPs showed high-confidence prediction results (i.e., with prediction AUC > 0.8) in the species closely related to human, including orangutan, monkey, mouse and rat (**Supplementary Fig. 7b**). In contrast, only a small fraction (32.9%–68.1%) of the RBPs showed high-confidence prediction results in the remaining distantly related species, such as yeast, *A. thaliana* and fly (**Supplementary Fig. 7b**). Taken together, these findings confirm that MuSIC performs significantly better in the closely related species, by leveraging the RBP conservation patterns in terms of both sequence and structure (**Supplementary Fig. 7b** and **Supplementary Table 5**).

### MuSIC improves RBP-binding motif identification in diverse species

Previous studies have revealed that conserved RBPs tend to have conserved RBP-binding motifs during evolution, which has been validated by some specific RBPs ^41,42^. However, this hypothesis has not been systematically tested in a broad set of RBPs. To this end, we first predicted the RBP-binding peaks across all the 11 species using the trained 2,046 RBP models, and identified the enriched RBP-binding motifs in the human and non-human species (**Supplementary Table 5**). The predicted enriched motifs in the non-human species could be successfully recapitulated by the well-known RBP binding sequence preference ^48^. For example, the motif CAUCU for YBX3, GGA for SRSF1 and polyC for KHSRP could be rediscovered in mouse, rat and chicken, respectively (**Fig. 4b** and **Supplementary Fig. 7c**). We then generated a catalog of RBP-binding motifs in all the 11 species ranging from human to yeast, and visualized the motif clustering based on their similarities (**Fig. 4c**; see **Methods** for details). Notably, RBP-binding motif prediction failed for certain RBPs due to their weak or undetectable binding signals in the distantly related species, such as yeast, fly, and zebrafish.

The inferred RBP-binding motifs with high confidence showed distinct degree of conservation during evolution (**Supplementary Fig. 7d** and **8**, **Supplementary Table 5**). For example, HNRNPA1 was predicted to contain GA-rich (e.g., GAA/AG) motifs in the closely related species (e.g., orangutan, monkey, and mouse) (**Supplementary Fig. 7d**). In contrast, the UGUANA-like motifs for PUM2 could be predicted in diverse species ranging from human to yeast (**Supplementary Fig. 7d**), consistent with the well-known motif UGUANAUA for PUM2 across multiple species ^51^. Overall, the predicted RBP-binding peaks by MuSIC enables the identification of RBP-binding motifs in diverse non-human species, revealing the conserved and diverged binding preference of specific RBPs.

### MuSIC reveals evolutionarily conserved RBP-binding peaks

It has been shown that the RBP-binding peaks from an RBP are either evolutionarily conserved or species-specific ^43,52^. The conservation patterns of RBP-binding peaks could be influenced by both the RBP itself and the conservation of the target RNA sequence. To systematically evaluate the conservation of RBP-binding across species, we analyzed the predicted RBP-binding peaks of 186 RBPs across 11 species in the conserved transcripts (**Supplementary Fig. 9a, b**; see **Methods** for details). As expected, we observed higher proportion of conserved RBP-binding peaks in the more closely related species (**Fig. 5a**). Almost 64% of the total RBPs analyzed here exhibited considerable global patterns of RBP-binding peaks conservation (**Fig. 5a**). We highlighted two RBPs, RBM15B and ALKBH1, with their RBP-binding peaks from 9 species on the evolutionarily conserved genomic region (**Fig. 5b**). Notably, there were fewer or none evolutionarily conserved RBP-binding peaks in the distantly related species.

**Figure 5:**
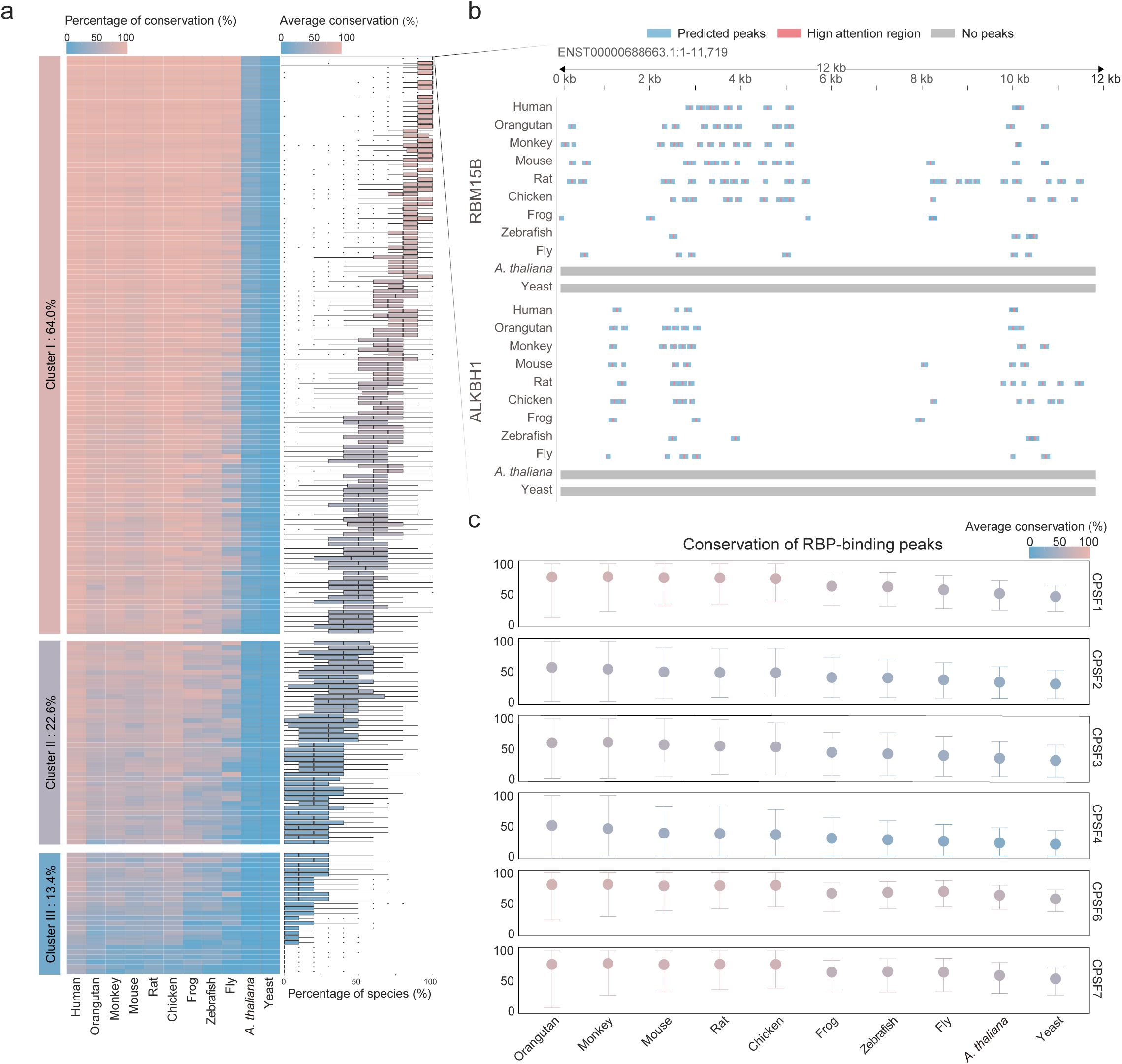
Cross-species conservation of RBP-binding peaks. (a) Heatmap showing the percentage of homologous transcripts containing the conserved peaks bound by RBPs across the 11 species, where all the 186 RBPs can be categorized into three groups. Boxplot showing the conservation degree (measured by percentage of species) of the homologous transcripts containing the conserved peaks. (b) Examples showing the predicted peaks bound by RBM15B and ALKBH1 along the transcript ENST00000688663.1. (c) Forest plot showing the conservation of the peaks bound by the CPSF family. We include the predicted peaks ranging from human to each of the non-human species shown here according to the phylogenetic tree. See also **Supplementary Fig. 9**.

We further investigated the conservation of RBP-binding peaks by analyzing the RBPs containing the same protein family from different species. The YTH family RBPs, which contain YTH protein domains, exhibited consistently strong RBP-binding peaks conservation across species (**Supplementary Fig. 9c**). In contrast, the CPSF family RBPs showed lower RBP-binding peaks conservation in the distantly related species (**Supplementary Fig. 9c**), consistent with previous studies reporting gradient conservation of the RBP-binding peaks ^43^. Further exploration of the relationship between RBP-binding peaks conservation and species evolution revealed that, while CPSF family RBPs maintained the steady degree of RBP-binding peaks conservation of in mammals, an obviously decrease showed among the distantly related species (i.e., from frog; **Fig. 5c**). Similar patterns could be also observed in the YTH family RBPs (**Supplementary Fig. 9d**). Taken together, these results indicate that the RNA-binding domains may play critical roles in maintaining the conserved RBP-binding peaks across species.

### MuSIC prioritizes genetic variation with consistent effects influencing RBP-binding across species

Genetic variations, such as SNVs, can potentially alter RBP-binding affinity by disrupting RNA target recognition, and thus cause functional abnormalities and severe diseases ^8,53,54^. The ability of predicting RBP-binding peaks across species by MuSIC enables us to quantify and analyze the effects of SNVs on RBP binding in the homologous regions between human and non-human species, which will be valuable for leveraging model organisms such as mouse to interpret the RBP-mediated mechanisms of human disease-associated SNVs ^10,55^.

We first quantified and compared the binding affinity on the local regions with (i.e., alternative allele) and without (i.e., reference allele) SNVs using the eCLIP data ^56^ from 103 RBPs (**Supplementary Fig. 10a**; see **Methods** for details). We found that SNVs detected in CLIP data were relevant to modulating RBP-binding affinity (**Supplementary Table 6**). Inspired by these findings, we further evaluated the consistency between MuSIC predictions and experimental data for SNV effects. We found that MuSIC could accurately predict the RBP-binding affinity on the local region containing the reference and alternative alleles, which was exemplified with the cases of SRSF1 and DDX3X (**Fig. 6a, b** and **Supplementary Fig. 10b**; see **Methods** for details).

**Figure 6:**
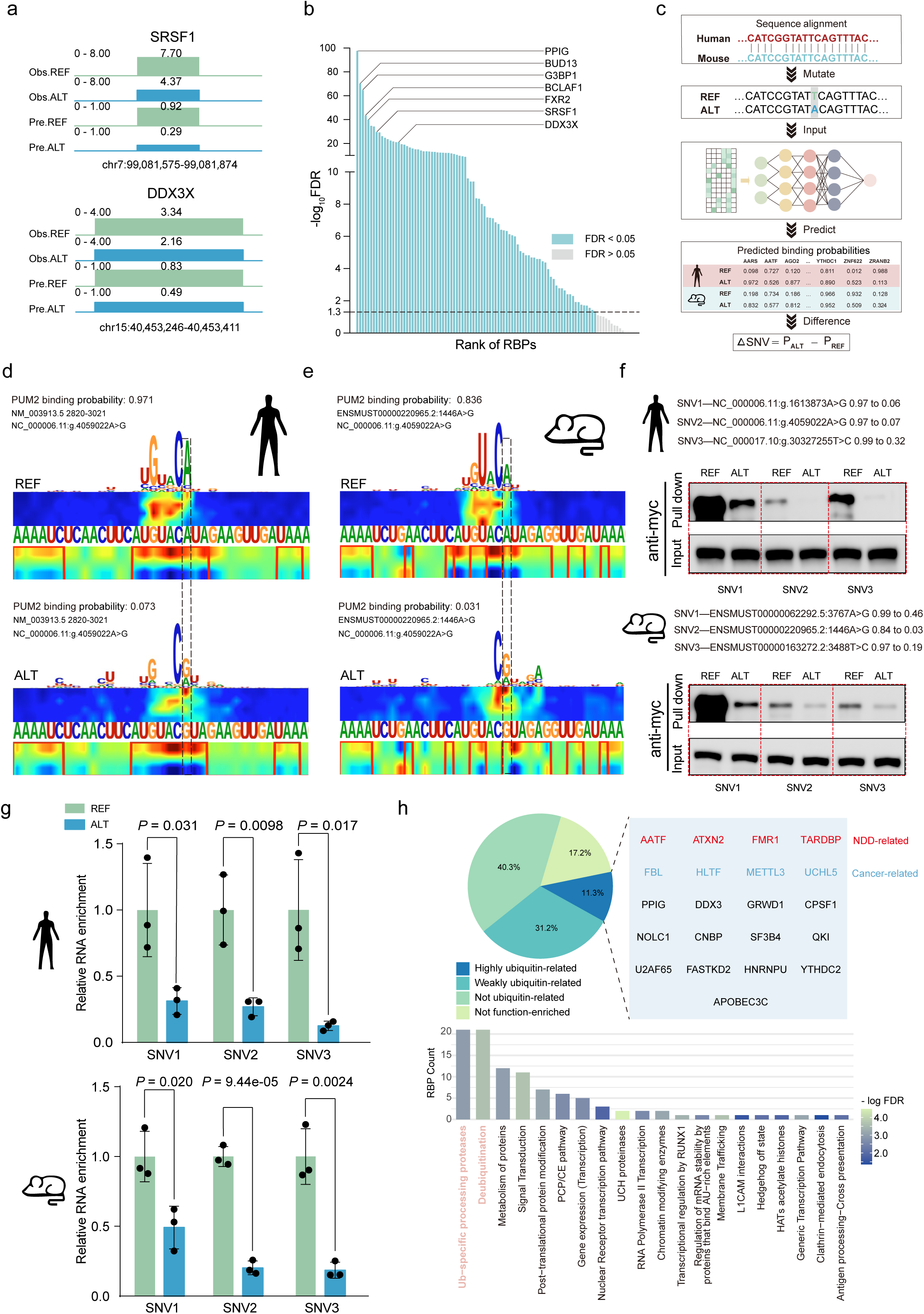
RBP–RNA interactions affected by SNVs and their associations with human diseases. (a) The experimentally-derived and predicted effects of SNVs on SRSF1 and DDX3X binding. Obs.REF, the binding strength from CLIP-seq data for the reference allele; Obs.ALT, the binding strength from CLIP-seq data for the alternative allele; Pre.REF, the binding strength from MuSIC prediction for the reference allele; and Pre.ALT, the binding strength from MuSIC prediction for the alternative allele. (b) Consistency between the experimentally-derived and predicted effects of SNVs on RBP binding for the 186 RBPs. Green bar indicates high consistency (-logFDR ≥ 1.3), whereas grey bar indicates no significance. (c) Schematic showing the workflow for quantifying the effects of homologous SNVs on RBP binding. (d) An example showing the effect of a homologous SNV (NC_000006.11: g.4059022A>G) on PUM2 binding in human. The saliency maps show the predicted effect of REF (A) (top) and ALT (G) (bottom) on PUM2 binding. In each saliency map, the two heatmap tracks display the sequence response (top) and structure response (bottom) of the model at the base level. The homologous SNV position is highlighted by a dashed box. (e) An example showing the effect of a homologous SNV (ENSMUST00000220965.2 1446 A>G) on PUM2 binding in mouse. The saliency maps show the predicted effect of REF (A) (top) and ALT (G) (bottom) on PUM2 binding. (f) Examples showing *in vitro* RNA pull-down assays (n = 3) examining the effects of three homologous SNVs on PUM2 binding in human (top) and mouse (bottom). (g) Examples showing *in vivo* POND-qPCR (n = 3) quantification of REF/ALT target RNAs binding to human and mouse PUM2 in HEK293T. Relative RNA enrichment was calculated as: (RNA ^supernatant^/ERCC spike-in)/ (RNA ^cellular^/GAPDH). Then the ALT group was normalized to the REF group. Green bar represents RNA enrichment for REF sequences, and blue bar for ALT sequences. (h) Enriched biological functions of the SNV-affected transcripts. The 186 RBPs can be categorized into four groups according to the enriched functions of their target transcripts. In total, 21 RBPs are identified as highly ubiquitin-related, some of which have been revealed as associated with neurodegenerative diseases (red) and cancer (blue). See also **Supplementary Fig. 10, 11, 12** and **13**.

Next, we ask whether MuSIC could be applied in quantifying the effects of SNVs in the homologous regions between species (**Fig. 6c**; see **Methods** for details). By applying the analysis framework, we systematically screened the 32,207 synonymous SNVs in human genome, corresponding to 85,599 SNVs in the homologous regions in mouse genome. We predicted 59,811 SNVs in mouse genome that could potentially disrupt RBP binding (i.e., ΔSNV < −0.5) (**Fig. 6c**). Here, we showed an example: the SNV NC_000006.11: g.4059022A>G in the human transcript NM_003913.5 was predicted as disrupting PUM2 binding (ΔSNV = −0.898) (**Fig. 6d**), consistent with the predicted effect of the SNV located in the homologous mouse transcript ENSMUST00000220965.2 (ΔSNV = −0.805) (**Fig. 6e**). Furthermore, we performed RNA pull-down *in vitro* and POND-qPCR ^57^ *in vivo* assays to validate the predictions by MuSIC. The POND (**<u>P</u>**rotein nanocage-emp**<u>O</u>**wered **<u>N</u>**on-**<u>D</u>**estructive) method is a recently developed novel technique for *in vivo* detection of RBP–RNA interactions, which enables selective packaging of RBP–RNA complexes into extracellular nanovesicles ^57^ (**Supplementary Fig. 11a**). We focused on three SNVs predicted by MuSIC showing disrupted binding affinity by PUM2 in both human and mouse (**Supplementary Table 7**). The RNA pull-down *in vitro* results confirmed that the alternative allele could indeed significantly impact PUM2 binding on the target RNA sequences in human and mouse (**Fig. 6f** and **Supplementary Fig. 11b, c**). In addition, the POND-qPCR *in vivo* results also supported these observations in human and mouse (**Fig. 6g**), confirming that MuSIC can generate accurate and reliable prediction on the effects of homologous SNVs on RBP binding across species.

### Pervasive influenced RBP-binding implicated in disease mechanisms

Among all the 186 RBPs analyzed for their binding affinity changes on the SNVs in mouse genome, 79 (42.5%) RBPs were found to be enriched with the biological processes associated with ubiquitination regulation and protein degradation (**Fig. 6h** and **Supplementary Fig. 12a, b**). Previous studies have revealed that ubiquitination-related processes are implicated in the pathogenesis of complex diseases, such as cancer ^58–61^ and neurodegenerative diseases ^62–65^, which can be also validated by the predictions by MuSIC (**Supplementary Fig. 12c**). These findings suggest the critical roles of SNVs in modulating RBP binding and influencing the ubiquitination-associated processes, which may be implicated in complex disease mechanisms through disrupting protein homeostasis.

In our analysis, TARDBP was prioritized as highly related to the ubiquitination-related processes. TARDBP has been reported to be critical in regulating protein homeostasis and protein folding, and abnormal TARDBP binding on the target RNA transcripts can lead to several diseases in human and mouse ^66–70^. We confirmed that the SNV in mouse could disrupt TARDBP binding to *Usp25*, a gene involved in ubiquitin metabolism (**Supplementary Fig. 13a**). Moreover, the TARDBP-bound genes perturbed by SNVs were significantly enriched in the functions related to neural development, ubiquitination, deubiquitination, and protein metabolism (**Supplementary Fig. 13b, c**).

## Discussion

Existing computational methods for predicting RBP–RNA interactions ^26,28,30,31^ were generally developed for the within-species tasks, having difficulties in predicting cross-species. In this study, we developed MuSIC, a novel computational method that can accurately predict RBP-binding peaks in the species lacking available CLIP datasets by leveraging the multi-level conservation patterns of the RBP. Compared to other similar computational methods, MuSIC showed superior performance in predicting RBP-binding peaks on the cross-species tasks. We then systematically predicted RBP-binding peaks and their enriched sequence motifs for 186 RBPs in human and 10 non-human species using MuSIC. We confirmed that the conservation patterns of the predicted RBP-binding peaks were consistent with previous studies ^42,43,52^. Finally, we applied MuSIC to investigate the effects of SNVs on RBP binding, particularly focusing on the imbalance in RBP binding induced by SNVs and exploring their potential connections to diseases such as cancer and neurodegenerative disorders.

The following improvements in the model architecture are critical for the augmented prediction performance of MuSIC in the cross-species tasks: optimization of input feature length and label smoothing. Previous studies have showed that increasing input sequence length can dramatically improve the model performance ^24,71^. In our study, we revealed that increasing the input sequence length from 101nt to 200nt could significantly improve the prediction performance across species (AUC from 0.55 to 0.75). In contrast, increasing the input sequence length from 200nt to 400nt would lead to decreased model performance. Increasing the input sequence length can not only provide more useful sequence information, but also may include noisy and redundant sequence information for modeling RBP binding. Inspired by the concepts from computer vision, we implemented label smoothing in the model to further improve prediction performance ^35–37^. We showed that label smoothing and the gradient weight adaptation mechanism could significantly improve cross-species predictions. This strategy reflects the inherently dynamic nature of biomolecular recognition, which operates on a continuum rather than a binary decision framework^72^.

The MuSIC model exhibited lower prediction confidence in the distantly related species, such as fly, *A. thaliana* and yeast, partially due to the decreased evolutionary conservation of RBPs in these species. With the advancement in large language models, RBP-guided RNA generation strategies could increase the prediction confidence of RBP-binding peaks in distantly related species ^73^. For example, BERT-RBP ^74^ was developed by adapting the pretrained BERT architecture and leveraging long-range RNA sequence features in predicting RNA-RBP interactions with improved performance. Similarly, ZHMolGraph ^75^ was proposed to predict novel RBP–RNA interactions by combining graph neural networks with unsupervised large language models. In the future, we hope that incorporating large language model-based diffusion methods into the MuSIC model can improve the prediction performance by complementing the low conservation of RBPs in the distantly related species.

Previous studies revealed that genetic variation in RNA can potentially reduce RBP-binding affinity ^8,53,76^. However, the effects of SNVs on RBP binding has not fully explored, particularly in the non-model organisms. In our study, we quantified the effects of SNVs on RBP binding in the homologous regions between human and mouse, and furthermore experimentally validated these prediction results *in vitro* and *in vivo*. Our results revealed the evolutionary conservation of genetic variation influencing RBP binding across species, and found that the SNVs that could potentially alter RBP–RNA interactions are associated with human diseases ^9,10^. We then performed a comprehensive analysis on the homologous human SNVs in mouse to quantify their binding affinity changes for each of the 186 RBPs. Interestingly, the disrupted binding events generally are recognized by TARDBP, and the target transcripts are enriched with the ubiquitination-associated pathways ^31^. The dysregulation of ubiquitin–proteasome system (UPS) can lead to protein misfolding and accumulation, which has been linked to human diseases including cancer and neurodegenerative disorders ^65,77^.

In summary, we propose MuSIC as a model to predict RBP-binding peaks across species. MuSIC outperformed state-of-the-art prediction models, and generated a catalog of predicted RBP-binding peaks of 186 RBPs with high accuracy in 11 species. The detailed analysis quantified the effects of genetic variants on RBP binding, particularly highlighted in the homologous regions between human and mouse. MuSIC provides insights into the RNA-RBP interactions in non-model organisms, and generates novel hypotheses on the interplays between RBP-mediated regulation and disease mechanisms.

## Supporting information

Supplementary Table 1: Evolutionary conservation of RBPs across 11 species

Supplementary Table 2: Conservation-based grouping of 216 RBPs

Supplementary Table 3: MuSIC training and validation datasets

Supplementary Table 4: Prediction performance of MuSIC

Supplementary Table 5: Predicted RBP-binding motifs from MuSIC

Supplementary Table 6: Consistency of SNV effects

Supplementary Table 7: Experimental validation of SNV effects

Supplementary Table 8: Functional enrichment of SNV-affected genes

## Acknowledgements

We thank Chenqian Wang, Shaozhen Yin, Ruobin Zhao, Liangyu Li, Yongkang Tang and Suiru Lu for fruitful discussions and invaluable feedback. We thank Jindong Sun for running the software comparison benchmark. This work was supported by the National Natural Science Foundation of China (No.82341086, No.32300521, and No.32422013 to L.S., No.32025007 to Y.W.); the Open Grant from the Pingyuan Laboratory (No.2023PY-OP-0104 to L.S.); the State Key Laboratory of Microbial Technology Open Projects Fund (No.M2023-20 to L.S.); the Intramural Joint Program Fund of the State Key Laboratory of Microbial Technology (NO.SKLMTIJP-2024-02 to L.S., and T.Z.); the Shandong Province Postdoctoral Innovation Project (NO. SDCX-ZG-202400146 to T.Z.), the Qingdao Postdoctoral Science Foundation (NO. QDBSH20240102200 to T.Z.), the Double-First Class Initiative of Shandong University School of Life Sciences; the Young Innovation Team of Shandong Higher Education Institutions, the Taishan Scholars Youth Expert Program of Shandong Province, and the Program of Shandong University Qilu Young Scholars.

## Author’s Contribution

L.S. conceived the project. L.S., Y.T.Y. and Y.W. supervised the project. J.H., and T.Z. established the initial technical framework. J.H. performed the computational analyses, data curation, and methods development. J.H., Q.C., S.Y. and W.Z. trained the deep learning models. L.H. and Y.W. conducted *in vivo* and *in vitro* experimental design and validation. Y.J., T.Z. and S.J. collected and re-analyzed the RBP similarity data. J.H. and J.W. collected and re-analyzed RBP-binding peaks data based on CLIP-seq. J.H., T.Z., and W.Z. deployed and tested the availability of the MuSIC scripts. J.H. and T.Z. written the initial draft of the manuscript. Y.T.Y., L.H., Q.C., J.Z., M.J., Y.L. and X.W. providing critical feedback and suggestions for improvement. J.H. formatted the figures, and L.S. and Y.T.Y. prepared the final version of the manuscript.

## Availability of data and materials

An implementation of the MuSIC script and benchmarking data is available at https://github.com/GALE1228/MuSIC_pretrain.

## Ethics approval and consent to participate

Not applicable.

## Consent for publication

Not applicable.

## Methods

### Compiling RBP-binding peak datasets

We obtained the RBP-binding peaks of 216 RBPs from POSTAR3 ^32^ for the within- and cross-species model training and testing. We applied a uniform preprocessing pipeline to the peak datasets to minimize putative bias in the datasets. Briefly, we first removed the duplicated and short (length < 5nt) peaks in each dataset. Then, we selected the top 5000 peaks with the highest binding strength in each dataset for the downstream analysis. Next, the peaks were extended by 200nt on both sides. We used GFF Annotation Parser (https://github.com/lipan6461188/GAP) to extract the RNA sequences of these extended peaks, which were considered as the positive peaks (i.e., RBP-binding peaks) in the model training and validation. We randomly selected 5000 RNA sequences of 200nt in length from the remaining genomics regions as the negative ones.

For the within-species prediction, 224 datasets of 186 RBPs from human, mouse, and zebrafish were used. For the cross-species prediction, 17 datasets of 13 RBPs shared between human and mouse, and 2 datasets of one RBP shared between human and zebrafish were used. Notably, the datasets of the same RBP generated by different CLIP-seq technologies were excluded in the cross-species analysis. For the within-species prediction, we used randomly 80% of the positive and negative peaks for the model training, whereas 20% were used for the model validation. For the cross-species prediction, we simply maintained a training-to-validation ratio of 4:1 by randomly selecting the positive and negative peaks from the source and target species.

### Reconstructing phylogenetic tree among the 11 species

We aim to develop a novel computational method to systematically predict RBP–RNA interactions in 11 species, including *Homo sapiens* (human), *Pongo abelii* (orangutan), *Macaca fascicularis* (monkey), *Mus musculus*(mouse), *Rattus norvegicus* (rat), *Gallus gallus* (chicken), *Xenopus laevis* (frog), *Danio rerio* (zebrafish), *Drosophila melanogaster* (fly), *Arabidopsis thaliana* (*A. thaliana*), and *Saccharomyces cerevisiae* (yeast).

We retrieved the 18S rRNA sequences of the 11 species from RNAcentral ^78^ (Release 24) for evaluating the evolutionary distance among these species. We then performed multiple sequence alignment of the 18S rRNA sequences using the Sequence Alignment tool in MEGA ^79^ v11.0.13, followed by constructing a phylogenetic tree among the 11 species using the neighbor-joining method. We then evaluated the reliability of the reconstructed phylogenetic tree using the Bootstrap test with 1000 replicates.

### Compiling sequence and structural datasets of RBPs

We collected the protein IDs of 216 RBPs ^32^ that were used in this study in the 11 species from UniProt ^51^ (**Supplementary Table 1**). For each RBP, the corresponding sequences and structural data were retrieved from UniProt ^51^. In the cases where predicted structural data for certain RBPs were unavailable in UniProt, we used AlphaFold2 ^80^ to predict and supplement the missing ones. The sequence similarity of the full-length RBPs between human and other species was computed using ClustalX^81^ v2.1.1. We used US-align ^82^ to evaluate the structural similarity of the full-length RBPs by calculating two metrics: TM-score (overall structure similarity) and pLDDT (confidence of predicted structure).

We obtained the annotations of 130 RBDs in the 11 species from InterPro ^83^ (Release 103.0), including their sequence positions and respective fragments in their full-length RBPs. The local structures of RBDs were extracted from the structures of full-length RBPs, ensuring the local structures of RBDs were aligned with the structures of full-length RBPs. Finally, ClustalX ^81^ v2.1.1 and US-align ^82^ were used to calculate the sequence and structural similarity of the RBDs between human and other species.

### Comparing RBP–RNA interactions in three-dimensional structures between species

We collected three-dimensional structures of homologous RBP RC3H1 (PDB ID 4QIL in human, PDB ID 4QI2 in mouse) and their bound RNA fragments from the PDB ^84^. In addition, we used AlphaFold3 ^49^ webserver to predict the interactions between homologous RBP IGF2BP1 and bound RNA fragments in human and mouse based on the IGF2BP1 protein sequence in human and mouse, as well as well-known RNA sequence with strong preference by IGF2BP1 (5’-UGACUUACUUCAUUAUUGAA-3’). The cartoon and surface diagrams displaying the bound RNA, RBDs, and the residues involved in the interactions were generated using PyMOL v3.1.0 (http://www.pymol.org/pymol).

### Pre-processing MuSIC input data

The RNA sequence of each peak in the datasets was first transformed into a one-hot encoded matrix of size (4, 200), where the four rows correspond to the nucleotides A, C, G, and U. The secondary structure of each RNA sequence was predicted using RNAfold ^85^ v2.1.0 with default settings. The resulting structure was encoded into a 2-dimensional matrix of size (2, 200), with ‘P’ and ‘U’ representing the paired and unpaired bases, respectively. The sequence and structure matrices were then concatenated along the channel axis to generate a unified (6, 200) representation encoding both sequence and structural features, which will be used as the input data for the MuSIC model.

### Overview of MuSIC architecture

The input data for MuSIC is defined as *S*, which contains *N* peaks. Each sample, *s* ∈ *S*, is an RNA sequence of length *L* = 200nt. As described above, *s* corresponds to a sequence-and-structure vector of D = 6 dimensions. Therefore, *S* is encoded into a tensor X ∈ *R*^*N*×D×L^ and the binding probability *Y* ∈ *R*^*N*^ is computed by as follows:

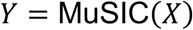

MuSIC is defined as follows:

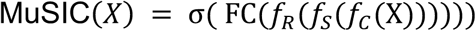

where *σ* is the sigmoid activation function, FC is the fully connected layer, *f*_*R*_ is the residual blocks, *f*_*S*_ is the squeeze-excitation block (SE), and *f*_*C*_ is the convolutional block, respectively.

### Convolutional Block

First, we use a convolutional layer to extract features. The convolutional block *f*_*C*_(X) is defined as follows:

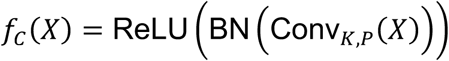

where Conv_*K*,*P*_(X) is a 2D convolutional layer with kernel K and padding P. The convolutional layer is designed to extract local sequence and structural features from the input data.

### Squeeze-and-Excitation (SE) Block

We apply a Squeeze-and-Excitation (SE) block to recalibrate channel-wise feature responses. The SE block is defined as follows:

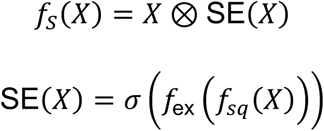

where *σ* is the activation function, *f*_*sq*_ is a global average pooling function to squeezes the global sequence and structural features into a channel statistic and *f*_ex_ applies a nonlinear transformation to the squeezed feature map. The SE block helps the model focus on the most relevant features for RBP binding.

### Residual Blocks

We then use residual blocks to capture both sequence and structural features over long ranges. These residual blocks are designed to learn the hierarchical features by passing information through several convolutional layers. Two types of residual blocks are used here:

ResidualBlock2D, which operates on 2D feature maps, is defined as follows:

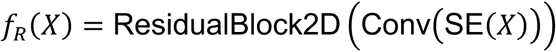

ResidualBlock1D, which operates on 1D feature maps, is defined as follows:

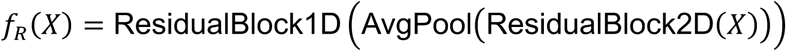

### Fully Connected Layer

Finally, we apply a fully connected layer to generate the binding probability prediction for each RNA input sequence, which is defined as follows:

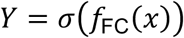

### Label smoothing strategy and gradient weight adaptation

We applied the label smoothing strategy to smooth the one-hot label distribution. Given a one-hot label vector *q*(*x*), the smoothed label distribution *q*’(*x*) is defined as follows:

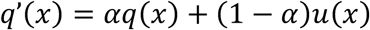

where *ɑ* is the RBP conservation score as a smoothing coefficient, and *u*(*x*) is the uniform distribution.

The one-hot label distribution *q*(*x*) and a predicted binding probability distribution *p*(*x*) are used for computing the classification loss as follows:

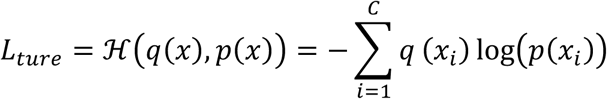

The smoothed label distribution *q*’(*x*) and a predicted binding probability distribution *p*(*x*) are used for computing the smoothing loss as follows:

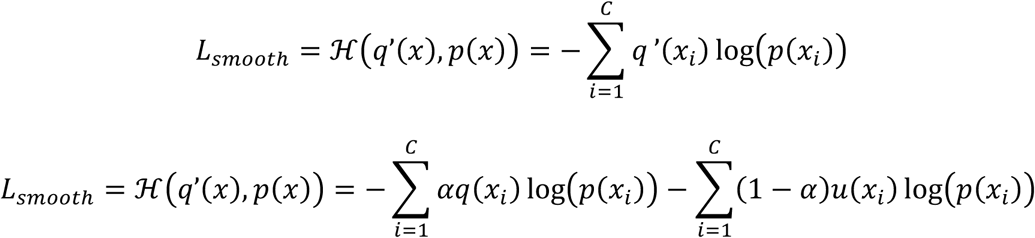

The first term, *L*_*true*_, represents the typical classification loss, while the second term, *L*_*smooth*_, is the smoothing loss. Finally, the total loss is defined as the weighted combination of classification loss and smoothing loss as follows:

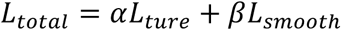

We proposed gradient weight adaptation to address the conflict between the gradient directions of *L*_*ture*_and *L*_*smooth*_. First, the gradients of both losses are computed with respect to the feature encoder parameters and classifier parameters as follows:

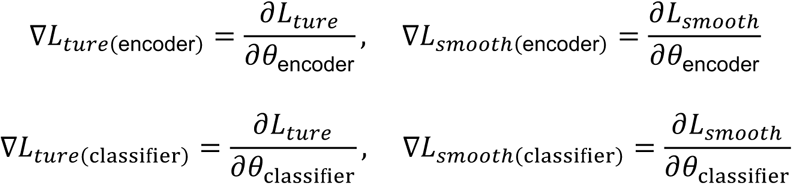

The weight of each loss term can be dynamically adjusted based on the ratio of the current loss to the previous one as follows:

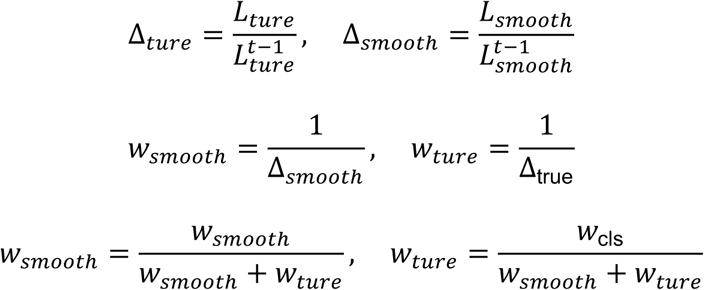

Subsequently, the gradient can be expressed as follows:

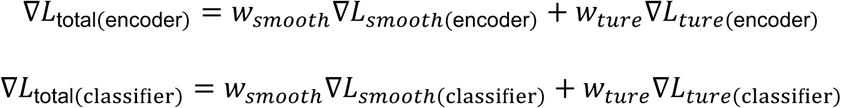

By dynamically adjusting *w*_*ture*_ and *w*_*smooth*_, the model can effectively prioritize the loss terms that show significant changes in the optimization process

### MuSIC model training

We applied the label smoothing strategy to smooth the one-hot label distribution and calculated the total loss as described above. The gradient weight adaptation dynamically modulated relative weights between *L*_*ture*_and *L*_*smooth*_according to the ratio of the current loss to the previous one, ensuring efficient gradient propagation.

The parameters in MuSIC model were optimized by minimizing the total loss— this loss integrates the classification loss and the smoothed label loss. We used Adam to optimize the model parameters, and Gradual Warmup Scheduler to adjust the learning rates. Throughout model training, we periodically evaluated the prediction performance of the model on the validation set to find the optimized parameters for the model.

## MuSIC hyperparameter settings

The hyperparameters of MuSIC were empirically adjusted based on the cross-species datasets. The hyperparameter settings for MuSIC are defined as follows:

Convolutional layer: the convolutional layer uses a kernel size of (3, 3) with 16 output channels, followed by Batch Normalization (BN) and ReLU activation for non-linearity.
Pooling layer: average pooling is applied with a kernel size of (6, 1), which reduces the feature map along the first axis of the input tensor.
Full connection layer: the fully connected layer at the end of the network has 1 output unit, which corresponds to the predicted binding probability for each sample.
Dropout probability: dropout is applied with probabilities of 0.1, 0.5, and 0.3 at different layers in the network to prevent overfitting.
L2 norm penalty: the L2 norm penalty (weight decay) is set to 1 × 10^-6^, which applies regularization to the model’s weights during optimization.
Batch size: the batch size is set to 64 for training and evaluation.
Learning rate: the learning rate is set to 0.001 for the Adam optimizer, which is used to optimize the network’s parameters.
Positive weight in loss function: the positive weight in the loss function is set to 2, which accounts for class imbalance in the dataset, giving more importance to positive samples during training.
Training epochs: the number of training epochs is set to 200, which determines how many times the model will iterate over the entire training dataset.
Early stop: early stopping is applied with a patience of 20 epochs, meaning training will halt if the model’s performance does not improve on the validation set for 20 consecutive epochs.
Initial weights: the initial weights of the model are set using the Kaiming normal initialization for convolutional layers and Xavier initialization for fully connected layers.

### Ablation experiments of MuSIC model

We performed the ablation experiments to explore to what degree the features could enhance the model performance: (i) input sequence length: we tried different input sequence lengths of 101nt, 200nt and 400nt; (ii) label smoothing strategy: we added and removed the label smoothing strategy in the models; (iii) sequence and structural features: we constructed the models using different types of input data, including only sequences, only predicted structures, combined sequences and predicted structures, and combined sequences and experimentally-measured structures.

### Performance comparison with other computational methods

We compared the prediction performance of MuSIC with other recently published computational methods, including HDRNet (https://github.com/zhuhr213/HDRNet) ^31^, PrismNet (https://github.com/kuixu/PrismNet) ^30^, DeepBind (https://github.com/jisraeli/DeepBind) ^28^, and GraphProt (https://github.com/dmaticzka/GraphProt) ^26^. Notably, to make a fair comparison, each of these computational methods was trained and validated on the same datasets as used in MuSIC, using their default parameter settings.

We extracted the features from the fully connected layers of MuSIC and the other four computational methods, including HDRNet ^31^, PrismNet ^30^, DeepBind ^28^, and GraphProt ^26^. These features were projected into a two-dimensional space using t-SNE to visualize and compare their classification performance.

We used MuSIC and the other four computational methods to train cross-species models and predict RBP-binding peaks for 22 RBPs from mouse and one RBP from zebrafish. We then estimated the prediction performance of the computational methods based on the overlap between the predicted peaks with binding probabilities above 0.9 and the experimentally-derived peaks. We also calculated the correlation between the predicted peaks and experimentally-derived peaks as we did in the meta-transcript analysis mentioned above.

### Thresholding high-confidence RBP across species predicted by MuSIC

We considered the six RBP features for RBP conservation analysis, including the full-length sequence similarity, the full-length structural TM-score, the full-length structural pLDDT, the RBD sequence similarity, the RBD structural TM-score, and the RBD structural pLDDT. We then applied Principal Component Analysis (PCA) to calculate the first principal component (PC1) as RBP conservation.

We then analyzed the correlation between the RBP conservation scores and the AUC scores of 14 RBPs derived from the cross-species analysis. We observed a linear correlation and constructed a regression model between the two variables, which can be used for estimating the cross-species prediction accuracy (i.e., AUC score) of the peaks (**Supplementary Table 4**). We applied a threshold of 0.8 for the AUC scores to retain the high-confidence predicted peaks.

### Identifying and clustering RBP-binding motifs

A total of 2046 models for predicting RBP–RNA interactions in 11 species were trained using 186 high-quality peak datasets from human ^32^. RNA sequences from each species were segmented into the fragments of 200nt in length, and 20,000 fragments were randomly sampled for further analysis. In each species, each of the MuSIC models of the 186 RBPs generated predicted peaks on these fragments. We then applied MEME ^86^ v5.5.7 to identify the enriched RBP-binding motifs using the fragments with binding probabilities above 0.9.

We performed motif clustering in two steps. First, for each motif, we calculated base frequency by averaging the nucleotide frequencies (A, C, G, U) from its position weight matrix (PWM) as follows:

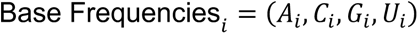

where *A*_*i*_, *C*_*i*_, *G*_*i*_, *U*_*i*_ are the average frequencies of adenine (A), cytosine (C), guanine (G), and uracil (U) for the motif *i*, respectively.

Second, we calculated the similarity matrix by integrating the Jensen–Shannon (JS) divergence of PWM distributions and the Euclidean distance between base frequency vectors:

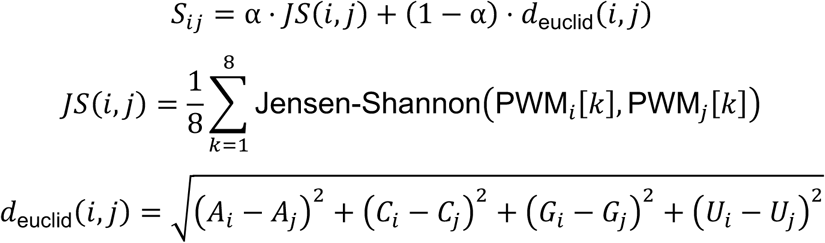

where PWM_*i*_[*k*] is the *k*-th row of the PWM matrix for motif *i*, PWM_j_[*k*] is the *k*-th row of the PWM matrix for motif *i*, *d*_euclid_(*i*, *j*) is the Euclidean distance between the base frequency vectors of motif *i* and *j*, *ɑ* is a hyperparameter balancing the weight of the JS divergence and Euclidean distance.

Finally, we grouped the motifs using hierarchical clustering to show their overall similarity. We used the itol.toolkit package ^87^ v1.1.10 to visualize and map the motif logo to the clustering results.

### Analyzing conservation patterns of RBP-binding peaks predicted by MuSIC

Using human transcripts as the reference, we applied the Mashmap ^88^ v3.1.3 to compare pairwise species (i.e., human and non-human species) and select the highest conserved transcripts with segment length > 500nt. For each species, MuSIC was used to predict RBP-binding peaks with 200nt and high attention regions ^30^ with 40nt for 186 RBPs in these conserved transcripts.

The conserved transcripts from 11 species were subjected to multi-sequence alignment by using Muscle5 ^89^ v5.1 with default parameters, and high attention region coordinates were aligned accordingly. RBP-binding peaks overlap counts across species were statistically analyzed within an extended window of 200nt. We then used IGV ^90^ v2.16.2 to visualize the RBP-binding peaks in the conserved transcripts from 11 species.

### Evaluating consistency between MuSIC predictions and experimental data for SNV effects

We downloaded the eCLIP peaks of 103 RBPs in HepG2 cell line from ENCODE (https://www.encodeproject.org/publication-data/ENCSR456FVU/). The SNVs in the RBP-binding peaks of each RBP were then identified from the eCLIP mapping files (i.e., BAM files) using the BCFtools mpileup ^91^ v1.20, where the replicates were merged. Peaks regions were filtered (p.adj ≤ 0.05) and merged across three replicates to retain high-confidence RBP-binding peaks. Using BEDTools ^92^ v2.27.1, SNV positions were overlapped with peak regions to classify peaks into SNV-overlapping and non-overlapping groups. A paired t-test was then applied to compare signal differences between the two groups.

We analyzed the RNA-seq alignment files from the smartSHAPE datasets (GSE145805) to estimate the background allele ratio. For both eCLIP and RNA-seq datasets, we calculated their reference allele (REF) and alternative allele (ALT) ratios using BCFtools mpileup ^91^ v1.20, and then obtained *diff*_*ratio*_, the difference between the REF and ALT ratios from the RNA-seq and eCLIP datasets as follows:

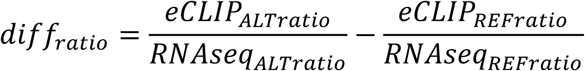

In parallel, we also calculated *diff*_*score*_, the predicted difference between the REF and ALT ratios as follows:

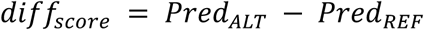

where *Pred*_*ALT*_ and *Pred*_*REF*_ are predicted binding possibilities for the sequences containing the ALT and REF, respectively.

We pre-processed the SNV effects by retaining |*diff*_*ratio*_| ≥ 0.2 and |*diff*_*score*_| ≥ 0.2, and then extracting the top 80% ranked by *diff*_*score*_. We used the experimental labels (*label*_*exp*_) and the predicted labels (*label*_*pre*_) to evaluate the consistency between experimental data and MuSIC predictions for SNV effects. The *label*_*exp*_ and *label*_*pre*_ are defined as follows:

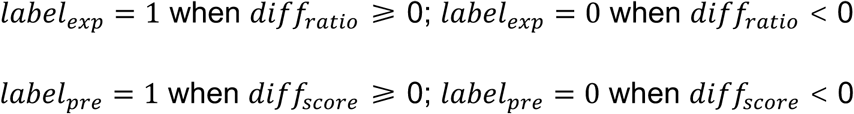

The consistency between *label*_*exp*_ and *label*_*pre*_ was evaluated using Fisher’s exact test, with multiple testing corrected by the Benjamini–Hochberg method, and results with FDR ≤ 0.05 were considered statistical significance.

### Quantifying the effects of homologous SNVs between human and mouse on RBP-binding using MuSIC

We downloaded the human and mouse RNA sequences from Ensembl ^93^ (Release 113). We extracted the human RNA sequences overlapping with the nonsense SNV positions from dbSNP ^6^ using the GAP (https://github.com/lipan6461188/GAP), and then generated the two sequences containing the REF and ALT for each SNV. We then identified the homologous regions of the human SNV-containing sequences in the mouse genome using BLAST ^94^ v2.16.0. In total, 32,207 nonsense SNVs in human, and 85,599 homologous nonsense SNVs in mouse were included for further analyses.

We then used MuSIC to predict the binding possibilities on the RNA sequences containing the REF and ALT in human and mouse for each of the 186 RBPs. For each sequence, the effect of the SNV on RBP binding was defined as follows:

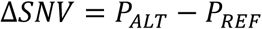

where *P*_*REF*_ is the binding probability for the REF, *P*_*ALT*_ is the binding probability for the ALT.

### Cell transfection

150,000 HEK293T cells per well were seeded in a well of poly-D-lysine-coated 24-well plates. After 16-20 h, cells were co-transfected with 200 ng EPN24-PUM2 plus 75ng GFP-REF or GFP-SNV target plasmid using JetPRIME reagent (Polyplus), the medium was refreshed at 4-6 h post-transfection, and the supernatant was collected for subsequent analysis 24 h after medium change.

### RNA pull-down assay

The *in vitro* RNA pull-down assay was performed as described ^95^. Briefly, 200 pmol biotinylated RNA oligonucleotides were refolded by heating at 90 °C for 1 min, and then incubated at 30 °C for 5 min. About 1 × 10^7^ myc-PUM2-expressing HEK293T cells were lysed in 500 µl lysis buffer (150 mM NaCl, 1 mM EDTA, 1% Triton X-100, 0.5 mM DTT, 50 mM Tris-HCl, pH 7.5, 0.5% sodium deoxycholate) with 5 µl PMSF (100mM, Beyotime), 10 µl phosphatase inhibitor cocktail (50 ×, Beyotime), 10 µl protease inhibitor cocktail (50 ×, Beyotime), 2.5 µl SUPERase In inhibitor (20U/µl Invitrogen) and 5 µl RNase Inhibitor (40U/µl, Beyotime). Refolded RNA (8 µl) and 500 µl cell lysate were incubated at 4 °C for 3-4 h. Then 100 µl pre-washed VAHTS CA-28 Streptavidin Beads (Vazyme) were added into the buffer and incubated at 4 °C for 1h. The beads were washed first with high salt buffer (50 mM Tris-HCl, pH 7.5, 1 M NaCl, 1% TRITON X-100) at room temperature for 5 min and then 2 more times with low salt buffer (50 mM Tris-HCl, pH 7.5, 150mM NaCl, 1% TRITON X-100), at room temperature and for 5 min. Proteins were eluted in 50 µl 1× SDS-PAGE loading buffer (Beyotime) at 98 °C for 20 min.

Protein electrophoresis was performed using BeyoGel™ Elite Precast PAGE Gel (8-16%, Beyotime), followed by protein transfer onto polyvinylidene fluoride (PVDF) membrane. The Myc Tag Mouse Monoclonal Antibody (HRP Conjugated) (Beyotime, AF2867) were diluted (1:1000) and incubated with the membrane at 4°C overnight. After three washes with 1×TBST buffer, the membrane was treated with electrochemiluminescence (ECL) reagent (Merck Millipore, WBKLS0050) and imaged with eBLOT14 Touch Imager (e-BLOT).

### POND-qPCR

Cell supernatant was clarified by centrifugation at 3,000 g for 5 minutes, then filtrated through a cellulose acetate filter with 0.45 μm pore size (Corning). 200 μl of supernatant was mixed with 800 μl TRIzol reagent (Magen) and 1 μl diluted External RNA Controls Consortium (ERCC, 1:1000) synthetic spike-in RNAs (ThermoFisher). RNA was extracted and dissolved in 20 μl nuclease-free water.

Next, 3 μl of supernatant RNAs or 250 ng cellular total RNA were reverse transcribed using HiScript® III RT Super Mix (Vazyme) at 37°C for 15 minutes. The cDNA products were diluted five-fold with 20 μl nuclease-free water, and 1 μl aliquot was used for real-time qPCR with AceQ qPCR SYBR Green Master Mix (Vazyme) in 96-well plates on the StepOne Plus Real-Time PCR System (Applied Biosystems) following standard protocols.

### Large-scale screening of RBPs affected by SNVs in UPS

For each RBP, SNVs with strong effects were identified: Δ*SNV* < −0.2 and *P*_*WT*_> 0.6. We annotated the transcripts affected by SNVs using the GAP (https://github.com/lipan6461188/GAP).

We used the STRING ^96^ database (https://string-db.org/) to annotate the enriched biological functions of SNV-affected transcripts. Among the top five enriched biological functions, we selected those related to ubiquitination or deubiquitination. Then, we categorized the RBPs into four groups according to the functional enrichment with ubiquitination or deubiquitination for their target transcripts affected by SNVs, i.e., the highly ubiquitin-related (both ubiquitination- and deubiquitination-related), the weakly ubiquitin-related (either ubiquitination- or deubiquitination-related), the not ubiquitin-related (not ubiquitination- or deubiquitination-related), and the not function-enriched.

We then performed the KEGG enrichment analysis on the SNV-affected transcripts annotated with highly ubiquitin-related. The enriched disease pathways revealed potential regulatory associations among SNVs, RBPs, and disease-relevant UPS genes.

### Quantification and statistical analysis

All statistical analyses were performed with R. Where applicable, the sample size (N) was specified either in the plot or in the corresponding figure legends. Analysis details could be found in the method section, such as statistical tests, definitions, etc. Boxplots were used to visualize the distribution of the data, with the box representing the interquartile range (IQR). The median was indicated by a horizontal line within the box, while the upper and lower hinges represented the 75^th^ and 25^th^ percentiles, respectively. The whiskers extended to the most extreme data points within 1.5 times the IQR from the box. Any data points beyond this range were considered outliers and plotted individually. Statistical significance was determined using appropriate tests, with *P* reported in the figure legends or main text.

## Supplementary Figure Legends

**Supplementary Figure 1:**
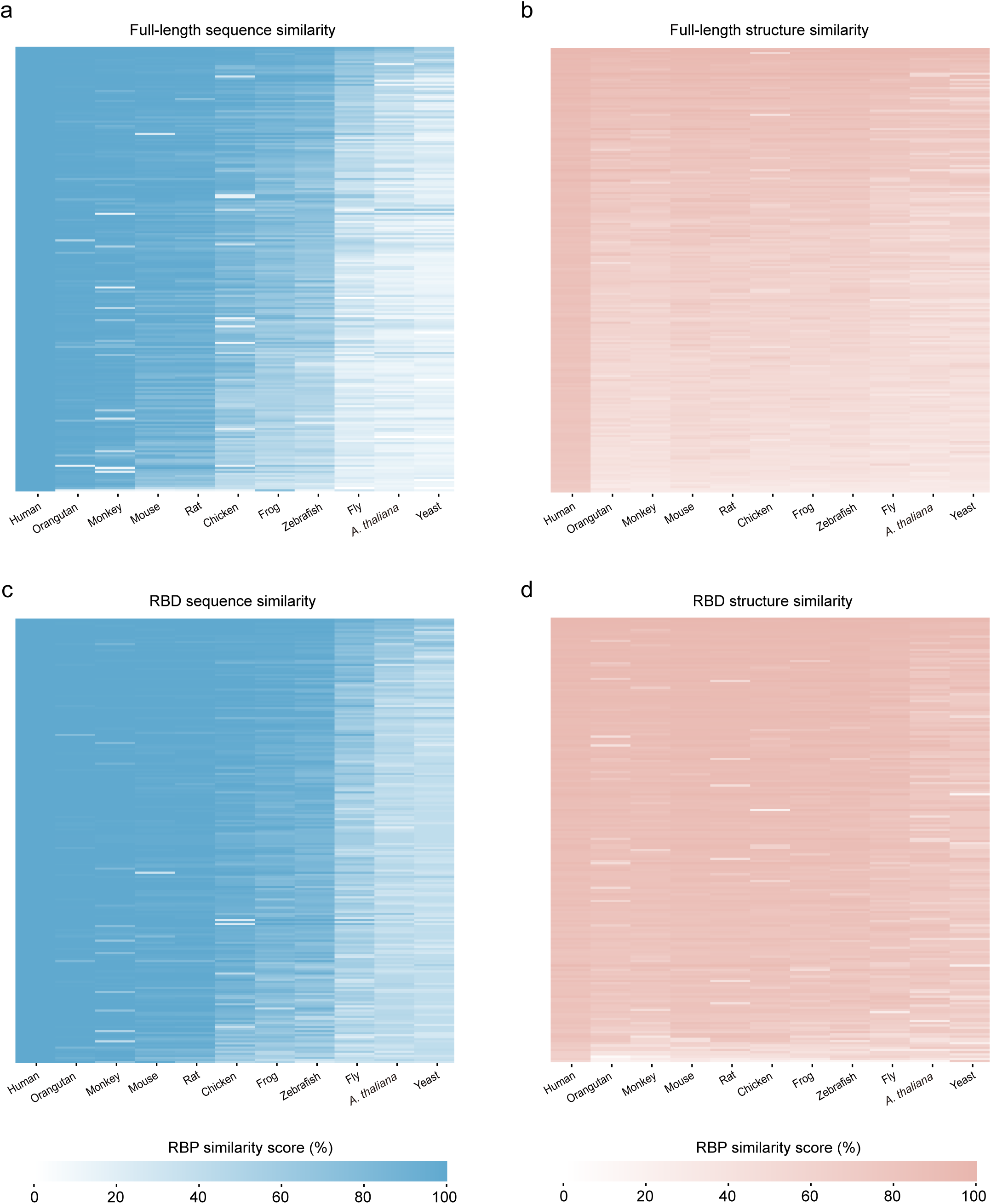
Evolutionary conservation of RBPs across 11 species. (a-b) Heatmaps showing the full-length sequence similarity (a) and structural similarity (b) of RBPs across species. (c-d) Heatmaps showing the RBD sequence similarity (c) and structural similarity (d) of RBPs across species

**Supplementary Figure 2:**
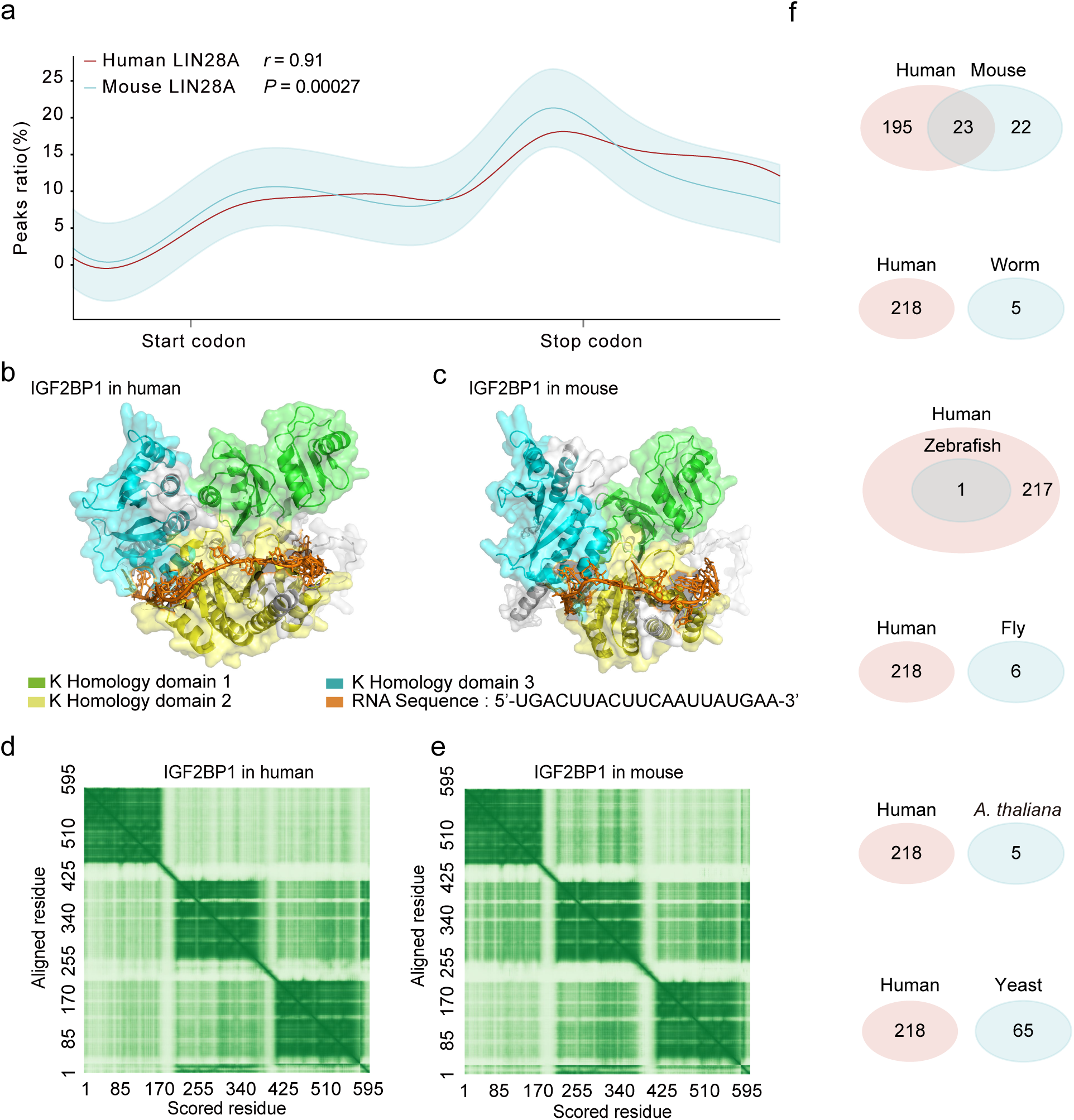
Evolutionary conservation of RBP-targeted RNAs. (a) LIN28A binding distribution along meta-transcript in human (red line) and mouse (cyan line). (b-c) Three-dimensional structures of IGF2BP1 in human (b) and mouse (c), showing the bound RNA residues and three KH domains residues. (d-e) Heatmaps showing residue interaction score for IGF2BP1 in human (d) and mouse (e), with darker green represents stronger interaction scores. (f) Venn diagrams showing the overlap of RBPs between different species (human, mouse, zebrafish, fly, worm, *A. thaliana*, and yeast) collected from POSTAR3 database.

**Supplementary Figure 3:**
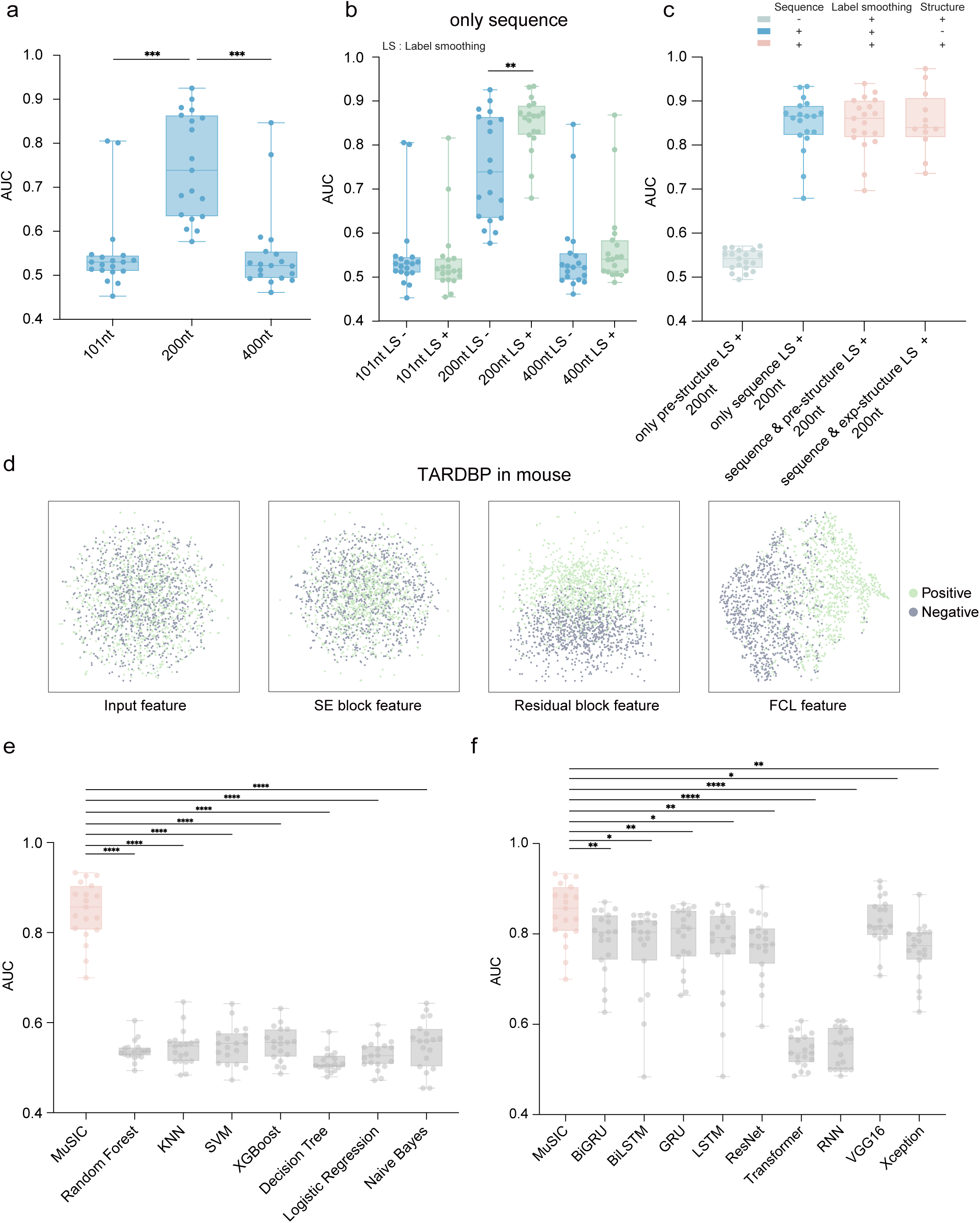
Optimization of input feature and model architecture. (a) Boxplot showing the prediction performance for different input sequence length (101nt, 200nt, and 400nt) on MuSIC performance. *** *P* < 0.005, paired t-test. (b) Boxplot showing the prediction performance of label smoothing, including only sequence (blue), and sequence with label smoothing (green). ** *P* < 0.01, paired t-test. (c) Boxplot showing the prediction performance of different structural features, including only predicted structure (cyan), only sequence (blue), sequence with predicted structure (red), and sequence with experimentally-measured structure (red). (d) t-SNE clustering showing the output feature maps by different blocks (input, SE block, residual block, FCL) for separating the positive (cyan) and negative (grey) peaks. (e) Boxplot showing the prediction accuracy of MuSIC (red) and other machine learning models (grey). **** *P* < 0.001, paired t-test. (f) Boxplot showing the prediction accuracy of MuSIC (red) and other deep learning models (grey). * *P* < 0.05, ** *P* < 0.01, **** *P* < 0.001, paired t-test.

**Supplementary Figure 4:**
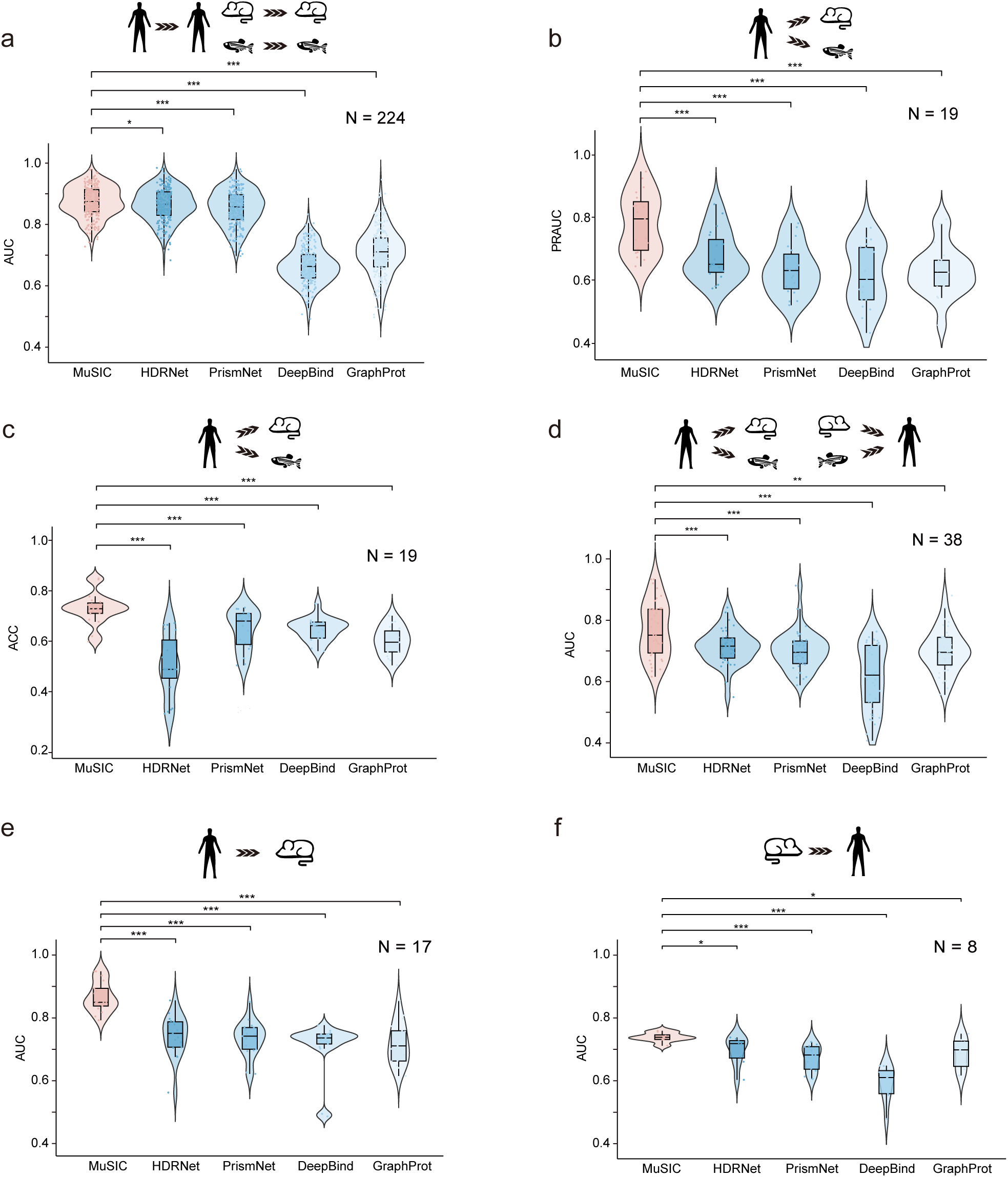
The performance comparison between MuSIC and other computational methods. (a) Violin plot showing the prediction accuracy of MuSIC (red) and other computational methods (blue) across 224 within-species datasets. * *P* < 0.05, *** *P* < 0.005, paired t-test. (b-c) Violin plot showing the prediction accuracy (PRAUC and ACC) of MuSIC (red) and other computational methods (blue) across 19 cross-species datasets. *** *P*< 0.005, paired t-test. (d) Violin plot showing the prediction accuracy of MuSIC (red) and other computational methods (blue) across 38 cross-species datasets. ** *P* < 0.01, *** *P* < 0.005, paired t-test. (e) Violin plot showing the prediction accuracy of MuSIC (red) and other computational methods (blue) across 17 cross-species datasets. *** *P* < 0.005, paired t-test. (f) Violin plot showing the prediction accuracy of MuSIC (red) and other computational methods (gradient blue) across 8 cross-species datasets. * *P* < 0.05, *** *P* < 0.005, paired t-test.

**Supplementary Figure 5:**
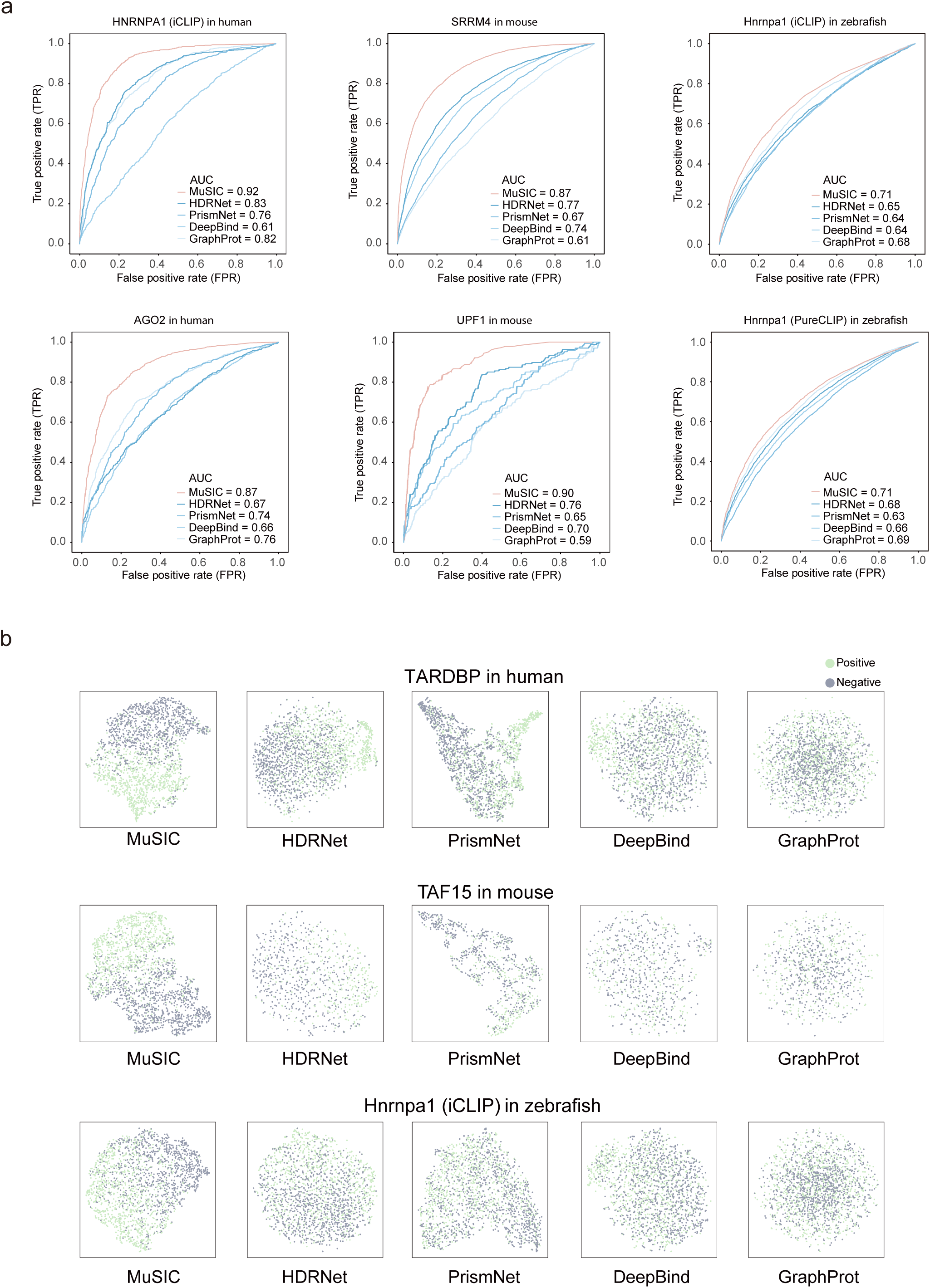
Comparison of feature learning discriminative power between MuSIC and other computational methods. (a) ROC curves showing the prediction accuracy of MuSIC (red) and other computational methods (blue) for 6 RBPs, 2RBPs for human (left), 2 RBPs for mouse (middle), and 2 RBPs for zebrafish (right). (b) t-SNE clustering showing the output feature maps by MuSIC and other computational methods for separating the positive (cyan) and negative (grey) peaks.

**Supplementary Figure 6:**
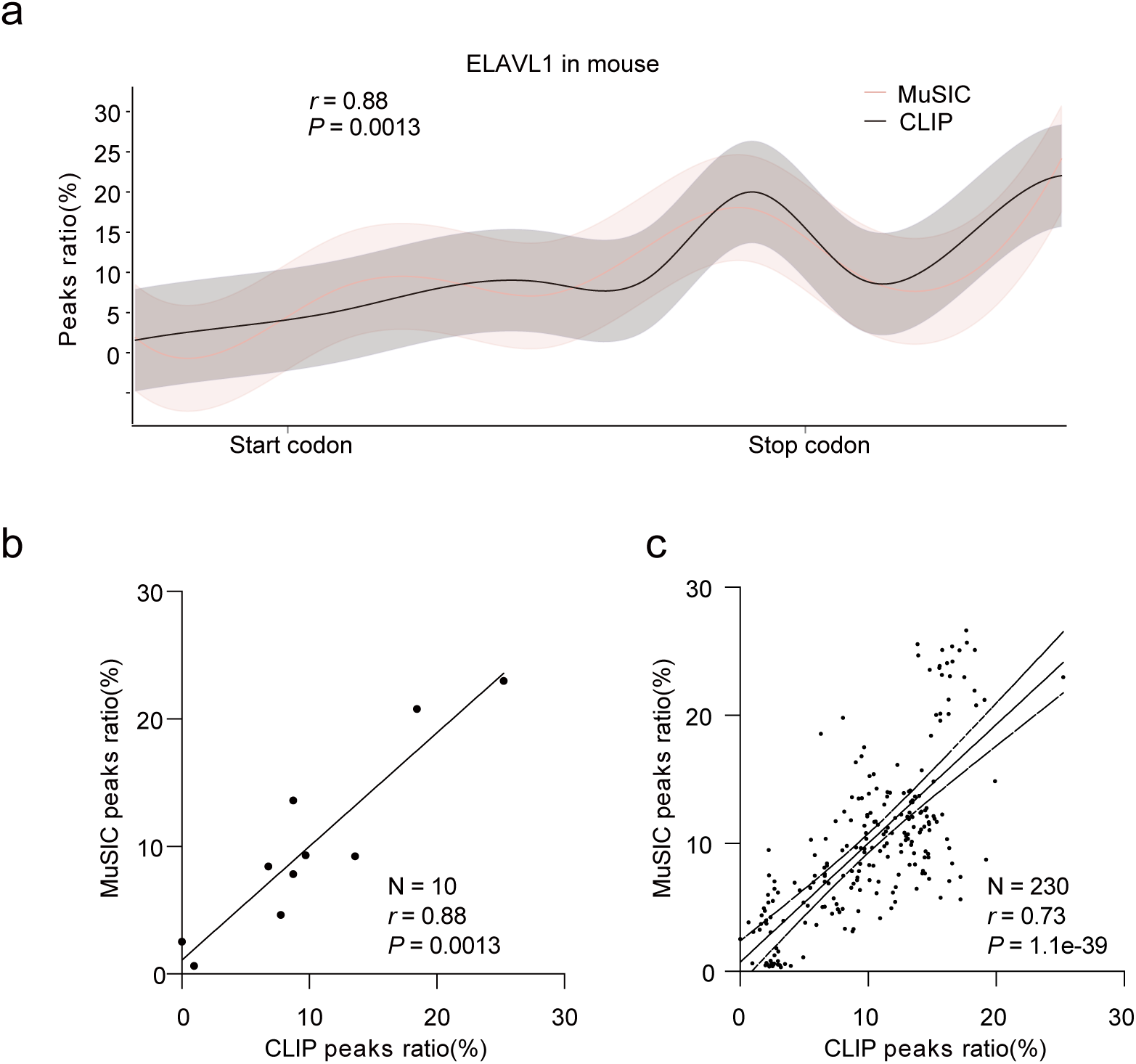
Consistency of RBP-binding distributions between MuSIC predictions and experimental data. (a-b) ELAVL1 binding distributions along meta-transcript in MuSIC predictions (red line) and experimentally-derived data (black line). Scatter plot showing the correlation of RBP-binding distributions from MuSIC predictions and experimental data for ELAVL1. (c) Scatter plot showing the correlation of RBP-binding distributions from MuSIC predictions and experimental data for 23 RBPs.

**Supplementary Figure 7:**
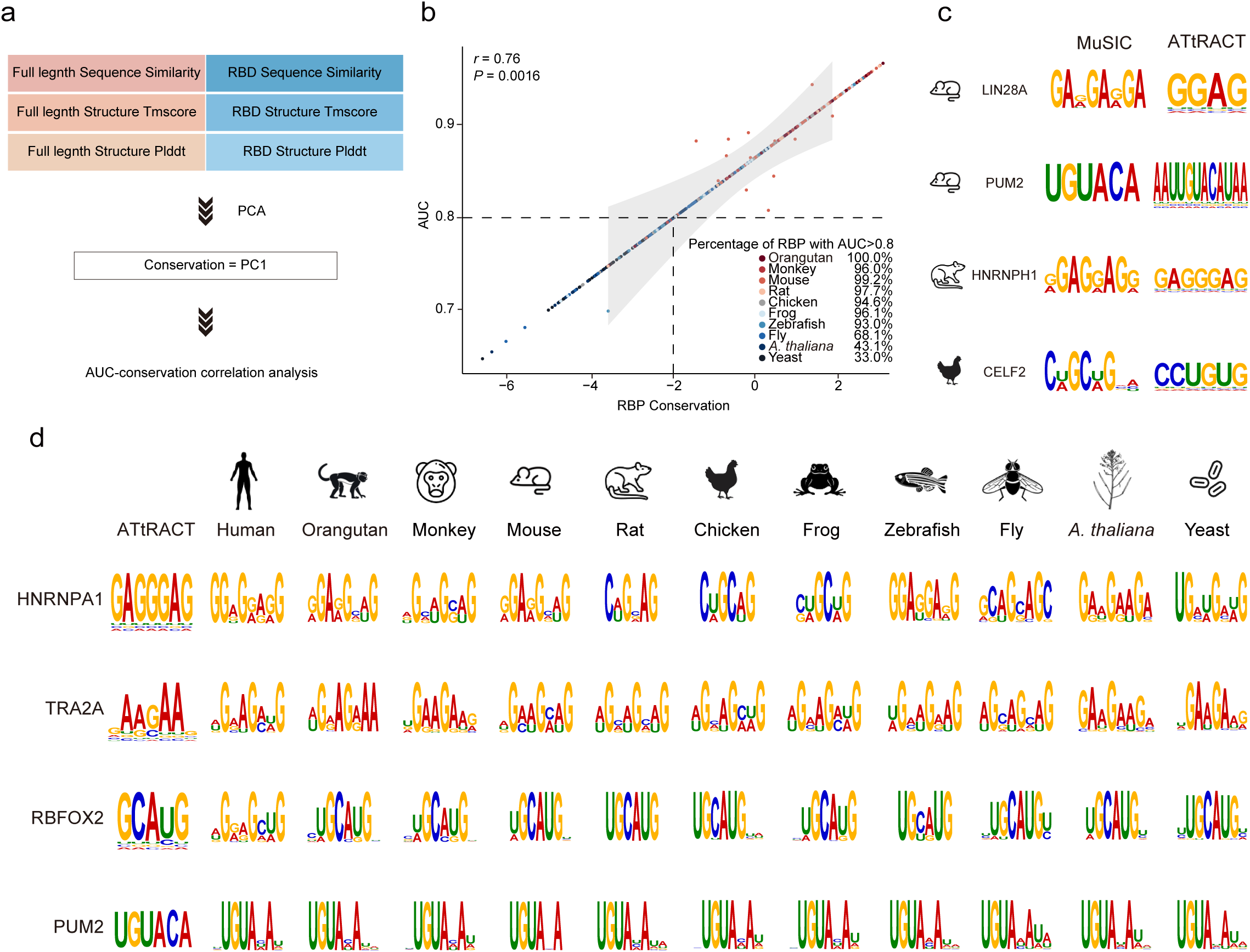
Evolutionary conservation of high-confidence RBP-binding motifs. (a) Flowchart showing the calculation of RBP conservation. (b) Scatter plot showing the correlation between the MuSIC-predicted accuracy and RBP conservation for RBPs from the 10 non-human species. (c) Examples showing the consistency of the predicted binding motifs and experimentally-derived motifs (from ATtRACT database) of several RBPs. (d) Examples showing high-confidence RBP-binding motifs across the 11 species, including human, orangutan, monkey, mouse, rat, chicken, frog, zebrafish, fly, *A. thaliana*, and yeast.

**Supplementary Figure 8:**
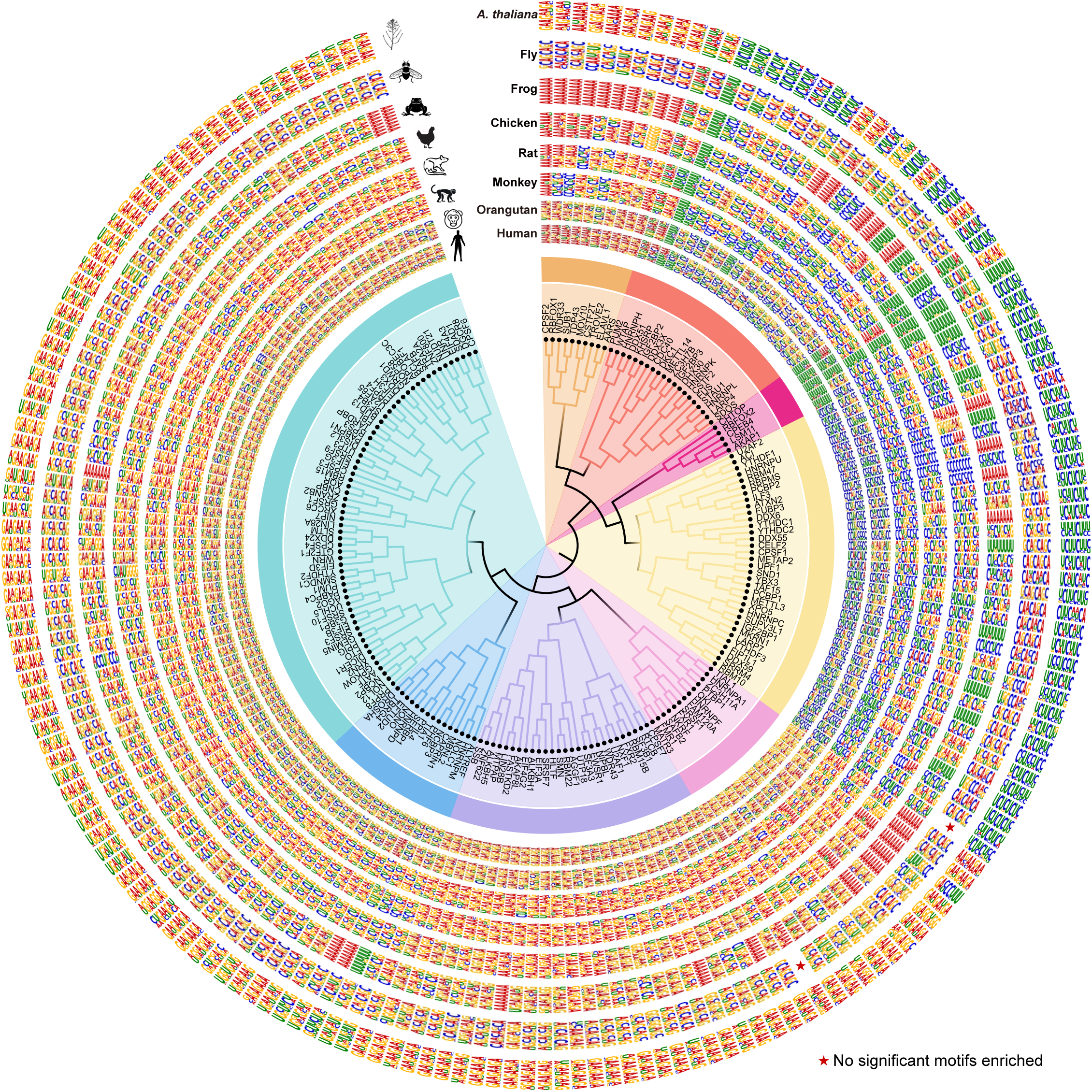
Clustering of RBP-binding motifs. Hierarchical clustering of 186 predicted binding motifs in human, orangutan, monkey, rat, chicken, frog, fly, and yeast. Stars indicate that there are no motifs enriched.

**Supplementary Figure 9:**
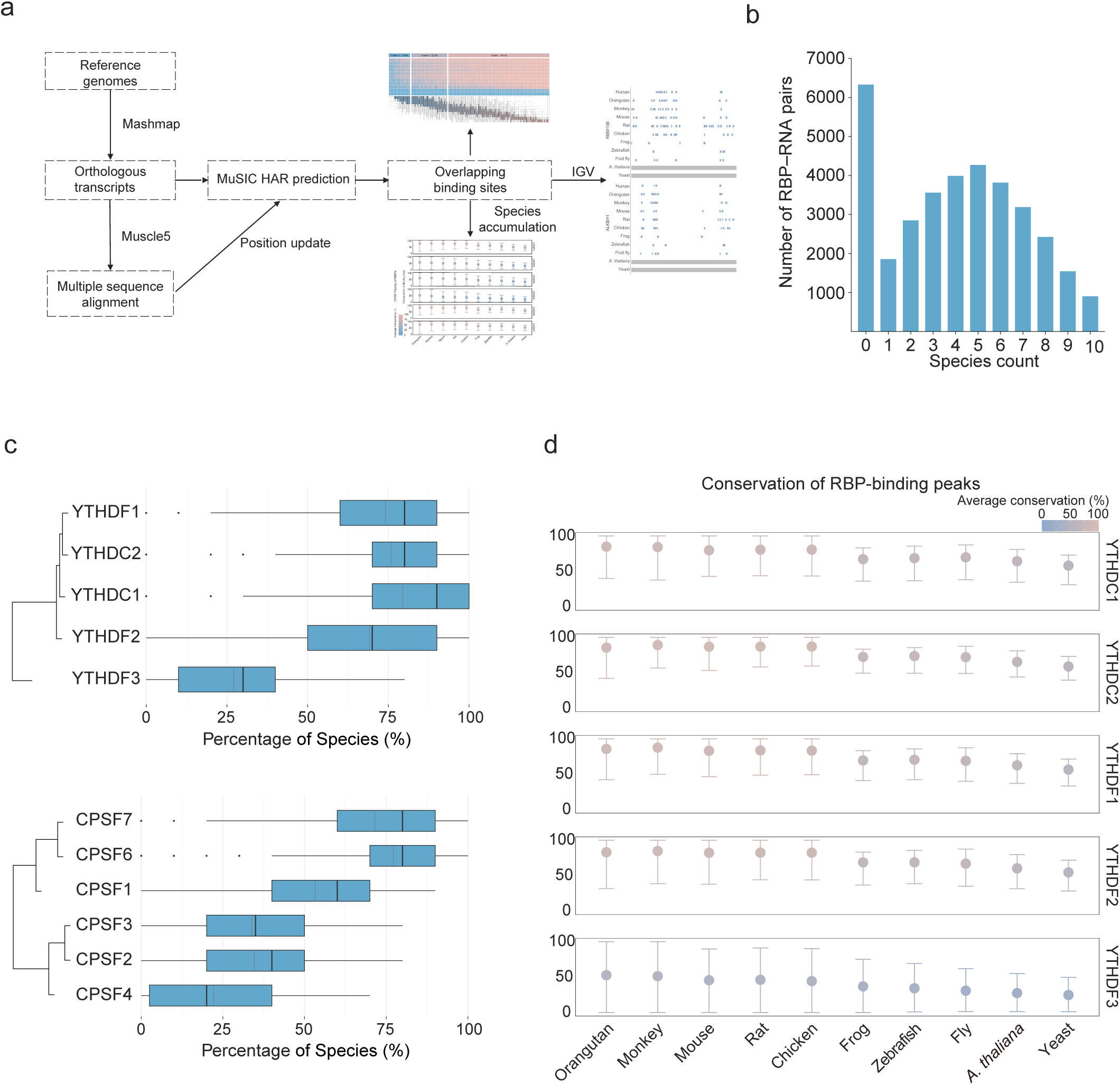
Cross-species conservation of predicted RBP-binding peaks. (a) Schematic showing the workflow for cross-species RBP-binding peaks conservation analysis. (b) Histogram showing the conservation of RBP-binding peaks across 11 species for 186 RBPs. (c) Boxplot showing the conservation degree of RBP-binding peaks across 11 species for CPSF family (top) and YTH (bottom) family. (d) Forest plot showing the conservation of the peaks bound by the YTH family. We include the predicted peaks ranging from human to each of the non-human species shown here according to the phylogenetic tree.

**Supplementary Figure 10:**
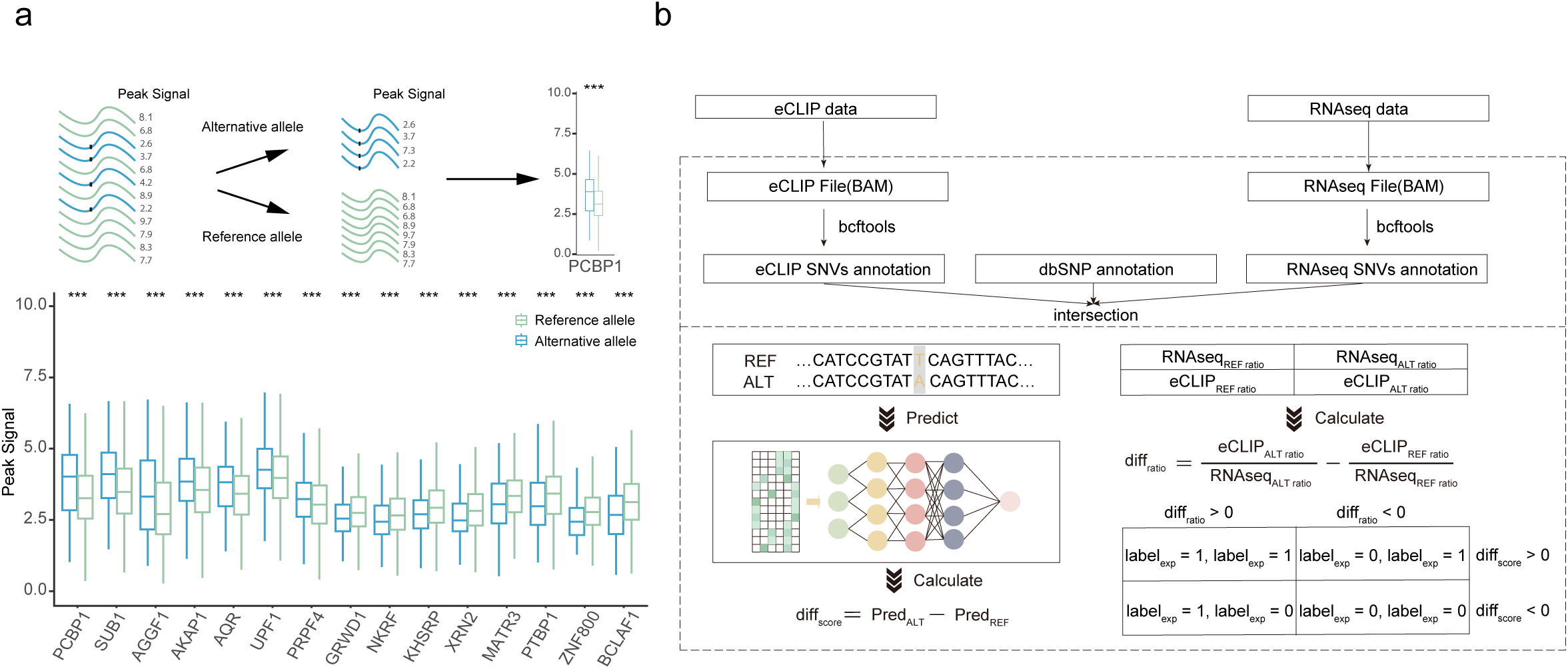
Consistency between the predicted and experimentally-derived effects of SNVs on RBP–RNA interactions. (a) Schematic showing the workflow for separating peak signals into the REF and ALT groups. *** *P* < 0.005, unpaired t-test. (b) Schematic showing the workflow for evaluating consistency of SNV effects between MuSIC predictions and experimental data.

**Supplementary Figure 11:**
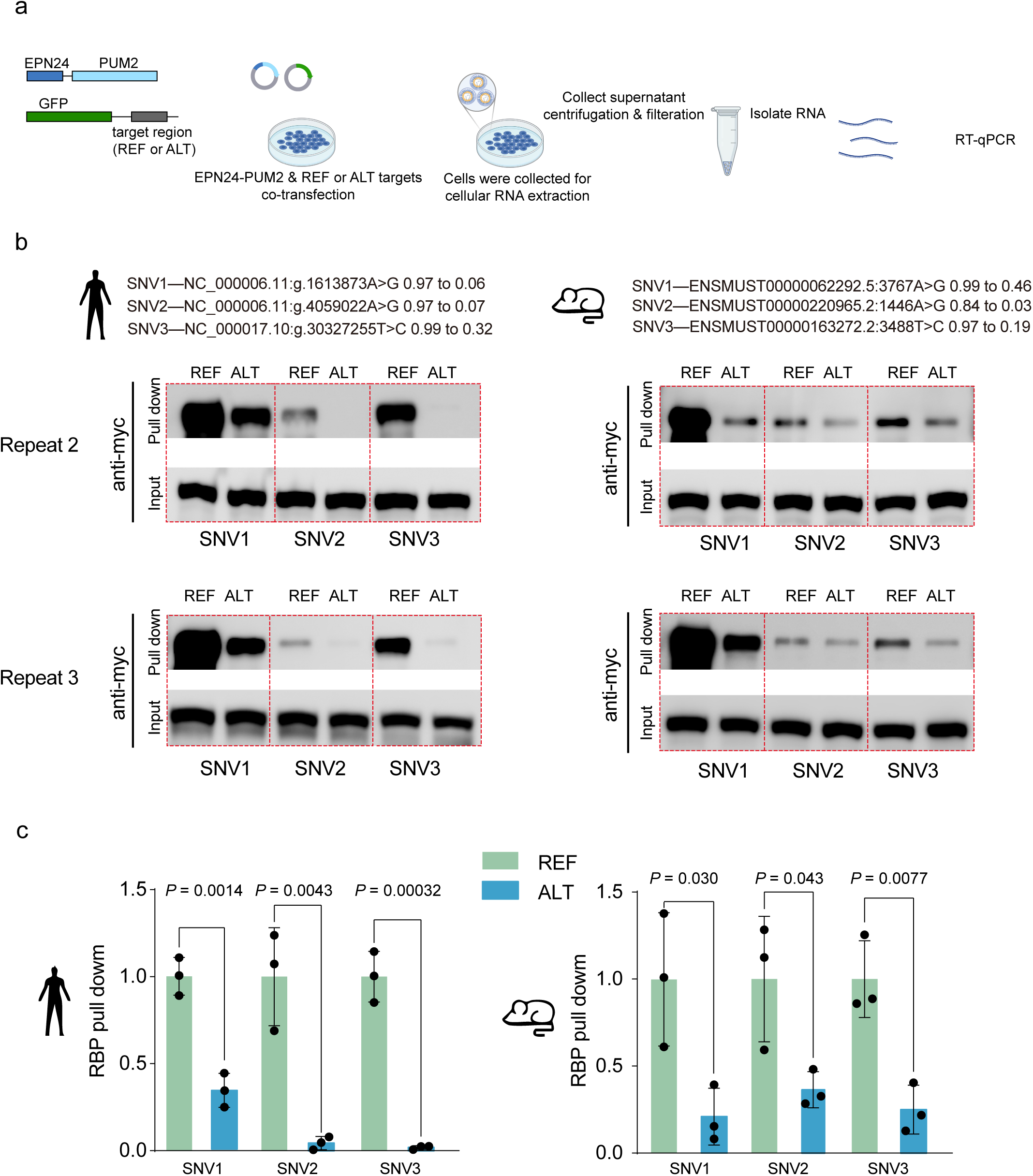
Experimental validations for the predicted effects of SNVs on RBP–RNA interactions in human and mouse. (a) Flowchart of using the POND method to detect the interaction between PUM2 and REF/ALT target RNA. PUM2 is fused to the C-terminus of the EPN24 nanocage monomer, while REF and ALT RNA targets are cloned into the 3’UTR of GFP on a separate plasmid. In HEK293T cells, EPN24-PUM2 is co-transfected with either GFP-REF or GFP-ALT RNA target plasmids. After transfection, EPN24-PUM2 packaged target RNAs into extracellular supernatant. Both supernatant and cells are collected for RNA extraction, followed by analysis of differential enrichment between REF and ALT target RNAs with RT-qPCR. (b) RNA pull-down assays and immunoblots for human PUM2 (left) and mouse PUM2 (right). Biotin-labeled REF and ALT RNA probes were used to perform RNA pull-down assays from lysates of 293T cells overexpressing myc-tagged PUM2. The pulled-down complexes were then analyzed by immunoblotting to detect the presence of PUM2 protein. The upper blot represents the results of repeat 2, and the lower blot represents the results of repeat 3. (c) Bar plot quantifying *in vitro* RNA pull-down (n = 3) enrichment of human and mouse PUM2 protein binding to REF and ALT target RNAs. For each condition, pulled-down PUM2 levels were normalized to PUM2 in the input lysate. Then the ALT group was normalized to the REF group. Green bars represent REF sequences, and blue bars represent ALT sequences. All *P* values are calculated using two-tailed unpaired Student’s t-test.

**Supplementary Figure 12:**
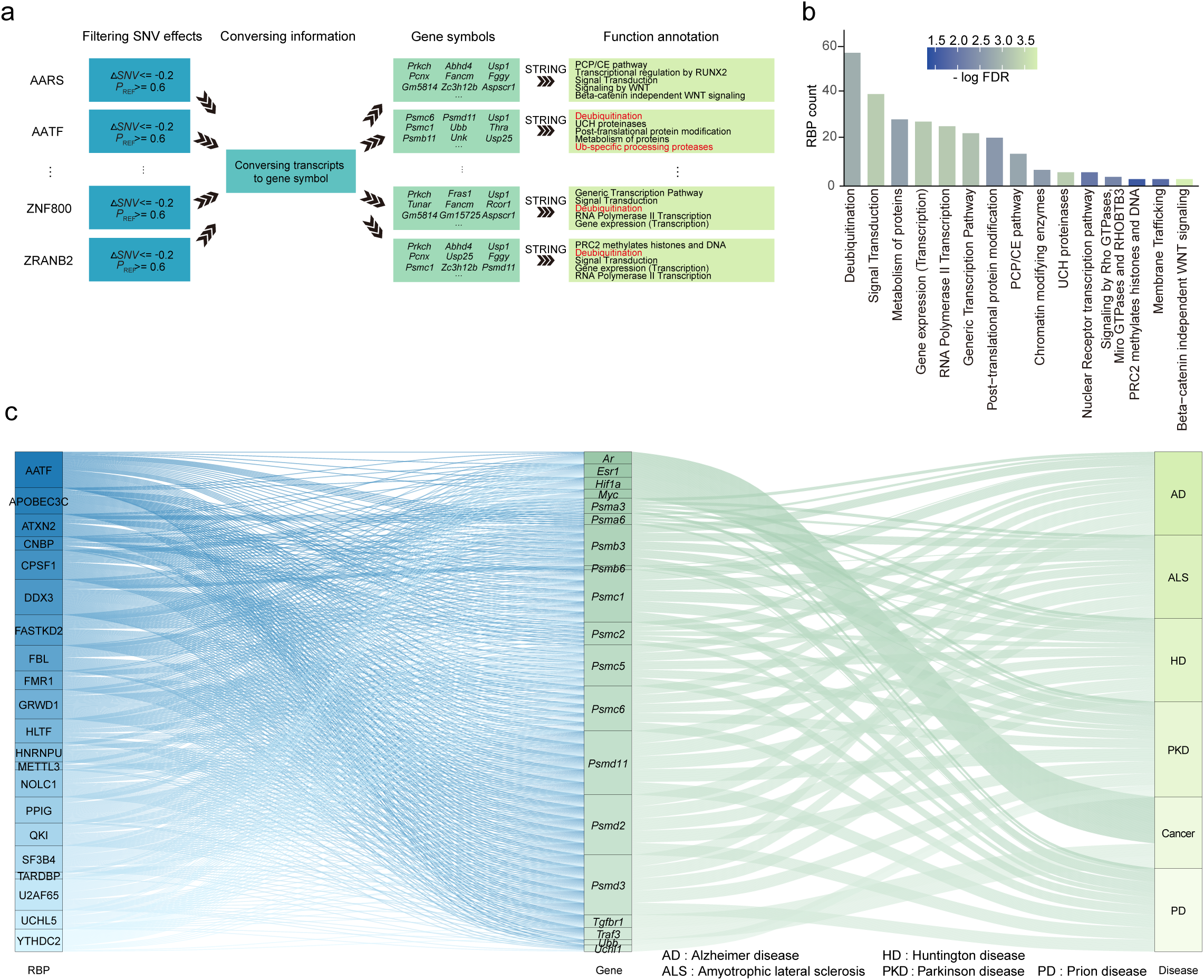
Predicted effects of SNVs on RBP–RNA interactions and their associations with human diseases. (a) Schematic showing the workflow of biological function enrichment analysis for RBPs affected by SNVs. (b) Bar chart showing the terms of biological functions associated with the weakly ubiquitin-related RBPs. (c) Sankey diagram showing the potential regulatory associations among SNVs, RBPs, and UPS-related disease.

**Supplementary Figure 13:**
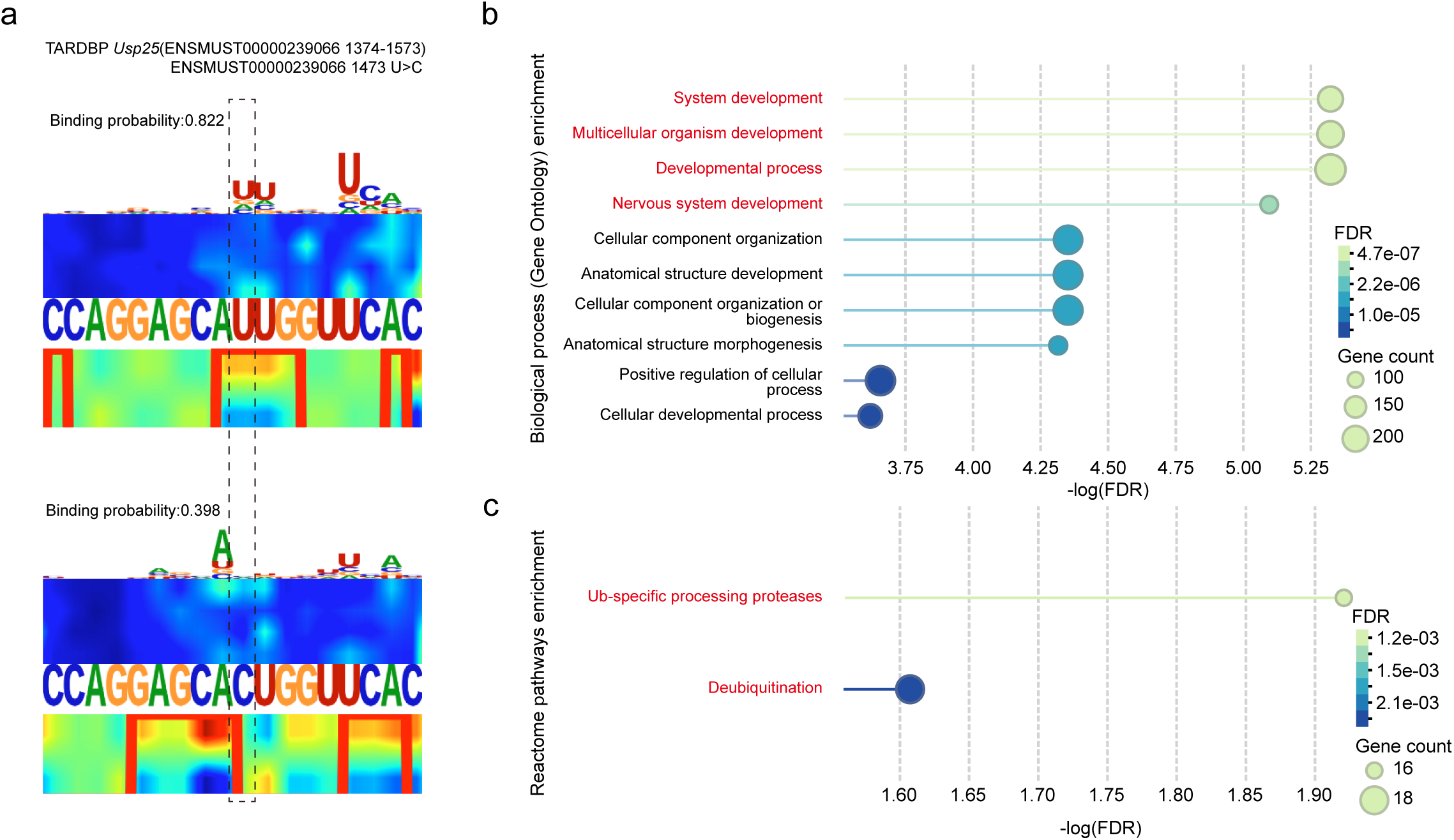
Predicted effects of SNVs on TARDBP binding and the enriched biological functions of the transcripts containing the SNVs. (a) An example showing the effect of a homologous SNV (ENSMUST00000239066 1473 U>C) on TARDBP binding in mouse. The saliency maps show the predicted effect of REF (U) (top) and ALT (C) (bottom) on TARDBP binding. (b) Gene Ontology enrichment result showing the biological process pathways for the TARDBP-bound genes perturbed by SNVs. (c) Reactome pathway enrichment result showing the biological functions for TARDBP-bound genes perturbed by SNVs.

## Supplementary Table Legends

**Supplementary Table 1: Evolutionary conservation of RBPs across 11 species**

**Supplementary Table 2: Conservation-based grouping of 216 RBPs**

**Supplementary Table 3: MuSIC training and validation datasets**

**Supplementary Table 4: Prediction performance of MuSIC**

**Supplementary Table 5: Predicted RBP-binding motifs from MuSIC**

**Supplementary Table 6: Consistency of SNV effects**

**Supplementary Table 7: Experimental validation of SNV effects**

**Supplementary Table 8: Functional enrichment of SNV-affected genes**

